# Chemokinesis by a microbial predator

**DOI:** 10.1101/2025.05.01.651543

**Authors:** Soniya R. Quick, Jason Bains, Catherine Gerdt, Bryan Walker, Eleanor B. Goldstone, Theresa Jakuszeit, Andrew W. Baggaley, Ottavio A. Croze, Joseph P. Gerdt

**Affiliations:** Department of Chemistry, Indiana University, Bloomington, IN 47405, USA; School of Mathematics, Statistics and Physics, Newcastle University, Newcastle upon Tyne NE1 7RU, United Kingdom; Institut Curie and Institut Pierre Gilles de Gennes, PSL Research University, CNRS UMR 144, 75005 Paris, France; Department of Mathematics and Statistics, Lancaster University, Lancaster, LA1 4YF, United Kingdom

## Abstract

Regulated motility is vital for many cells—both for unicellular microbes and for cells within multicellular bodies. Different conditions require different rates and directions of movement. For the microbial predator *Capsaspora owczarzaki*, its motility is likely essential for predation. This organism has been shown to prey on diverse organisms, including the schistosome parasites that co-reside with it in *Biomphalaria glabrata* snails. *Capsaspora* is also an evolutionary model for the unicellular ancestor of animals. This phylogenic placement makes *Capsaspora*’s motility an attractive target for understanding the evolution of motility in animal cells. Until now, little was known of how *Capsaspora* regulates it rate and direction of motility. Here we found that it exhibits chemokinesis (increased movement in response to chemical factors) in response to proteins released from prey cells. Chemokinesis also occurs in response to pure proteins—including bovine serum albumin. We found that this chemokinesis behavior is dependent on *Capsaspora* cell density, which suggests that the regulated motility is a cooperative behavior (possibly to improve cooperative feeding). We developed a mathematical model of *Capsaspora* motility and found that chemokinesis alone does not benefit *Capsaspora* predation. However, when coupled with chemotaxis (directional motility along a chemical gradient toward prey), chemokinesis may improve predation. Finally, we quantitatively analyzed *Capsaspora*’s previously reported chemotaxis behavior. These findings lay a foundation for characterizing the mechanisms of regulated motility in a predator of a human pathogen and a model for the ancestor of animals.

## INTRODUCTION

Motility is a key trait of many cells across the kingdoms of life. Motility allows unicellular organisms to enter environments with more nutrients, evade predators and toxins, and find partners for cooperative behaviors.^1,2^ Within multicellular organisms, motile cell types are essential for tissue development, immunity, wound healing, and sexual reproduction.^3^ In many cases, motility is tightly regulated. First, cells can regulate *when* they move (and relatedly, how fast they should move). Second, cells can regulate the *direction* in which they move. When the rate of movement is regulated by soluble chemical factors, the process is called chemokinesis; whereas, when the direction of movement is regulated by a gradient of soluble chemical factors, the process is called chemotaxis (**Figure 1A**).^4^ The molecular mechanisms of chemotaxis and chemokinesis are sparsely elucidated across different lineages, leaving open the question of how evolutionarily conserved these common behaviors are. Furthermore, the ecological implications of chemokinesis are still incompletely understood.^5,6^

**Figure 1.**
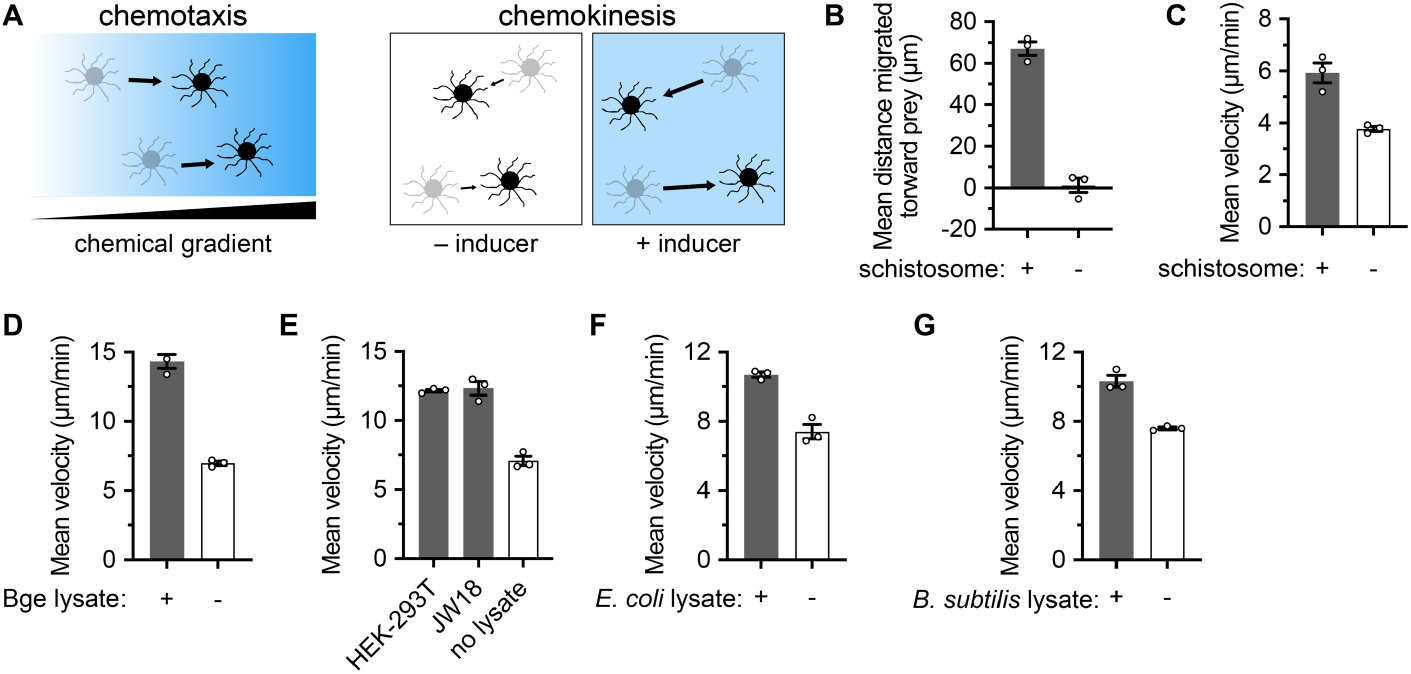
*Capsaspora* chemotaxis and chemokinesis responses to prey components. (A) Illustrations of chemotaxis (directional motility along a concentration gradient of a chemoattractant) and chemokinesis (increased rate of cell movement in response to a chemical inducer, without directionality). (B) Chemotaxis of *Capsaspora* toward schistosome sporocysts. Average net movement toward the schistosome is reported (negative value would indicate movement away from schistosome). Data were derived from **Videos S1-S3** (no schistosome) and **Videos S4-S6** (with schistosome). Only *Capsaspora* cells 70 μm to 240 μm from the schistosome were tracked. Data for individual cell tracks are reported in **Figure S1**. (C) Chemokinesis of *Capsaspora* in response to schistosome sporocysts. Non-directional cell velocity is reported, using cells tracked from the same videos for panel B. (D) Chemokinesis in response to lysate from *B. glabrata* embryonic (Bge) cells. Lysate from ∼2 × 10^6^ cells was added to 100 μl of *Capsaspora* culture. See **Figure S2A** for full dose-response data. (G) Chemokinesis in response to lysate from mammalian (HEK-293T) and insect (JW18) cells. Lysates from ∼2 × 10^5^ HEK-293T cells and ∼4 × 10^5^ JW18 cells were each added to 100 μl of *Capsaspora* culture. See **Figure S2B–C** for full dose-response data. (H) Chemokinesis in response to lysate from *E. coli* cells. Lysate from approximately ∼3 × 10^8^ cells was added to 100 μl of *Capsaspora* culture. See **Figure S2D** for full dose-response data. (I) Chemokinesis in response to lysate from *B. subtilis* cells. Lysate from ∼2 × 10^7^ cells was added to 100 μl of *Capsaspora* culture. See **Figure S2E** for full dose-response data. In all plots, error bars represent standard error of the mean of a biological triplicate of three individual wells of cells (n=3). Individual biological replicates are displayed with white circles. Within each biological replicate, dozens of individual cells were tracked. Chemokinesis assay method B was used for panels D–G (see **Methods**).

In order to shed light on the evolution of and ecological benefits of cellular motility, we here study the model protist *Capsaspora owczarzaki* (hereafter *Capsaspora*). *Capsaspora*’s multicellular aggregation behavior has been well studied;^7-12^ however, it also migrates in a mesenchymal-like fashion^13^ employing its long actin-filled filopodia.^14^ This motility behavior is notably less understood than (but just as vital as) the aggregation phenotype. *Capsaspora* was initially isolated from a *Biomphalaria glabrata* snail—a vector that transmits parasitic worms that cause schistosomiasis.^15^ In that initial report, *Capsaspora* was said to exhibit chemotaxis toward leaking schistosome prey. This behavior is proposed to be important for its ability to ‘hunt’ and devour schistosomes—which could enable *Capsaspora* to clear schistosomes from their snail vectors before they transmit to infect humans. *Capsaspora* is also one of the closest living relatives of multicellular animals. It shares many of the cell signaling and adhesion genes that are key for animal multicellular behaviors.^16^ Therefore, it is plausible that *Capsaspora* regulates its motility in ways that are uniquely homologous to animals—which would reveal the evolutionary timeline of cellular migration mechanisms in the animal lineage. Therefore, to determine both the ecological impact of *Capsaspora* motility and the possible ancestry of animal cell motility, we aimed to characterize the regulated motility of *Capsaspora*.

Here, we discovered that *Capsaspora* not only chemotaxes toward leaking schistosomes (as reported before^15^), but it also exhibits increased motility (chemokinesis) in response to leaking schistosomes. Furthermore, this chemokinesis response is promiscuous to lysates of diverse cell types, even including prokaryotes. We found discrete pure proteins that can trigger *Capsaspora* chemokinesis. It is common for cells to desensitize to stimuli over time,^17,18^ and we found that to be true here with *Capsaspora* chemokinesis, as well. Intriguingly, the chemokinesis response is dependent on the cell density of *Capsaspora*, but not due to direct cell-cell contacts. This may arise from cell-cell signaling through soluble chemical signals produced by the *Capsaspora* cells. Finally, to determine the benefit of chemokinesis for *Capsaspora*, we modeled its chemokinesis response and assessed if chemokinesis alone could improve *Capsaspora*’s ability to encounter its prey. This model showed no major advantage to chemokinesis alone, suggesting that chemokinesis synergizes with chemotaxis to increase the rate of directional motility for the predator’s benefit— as has been shown in a bacterial system.^5^ Furthermore, a slight benefit may exist for cells to disperse away from depleted prey. In total, this work provides initial mechanistic and ecological insight into regulated motility in a protist that may prevent the spread of schistosomiasis and may reveal the origins of cellular behaviors in multicellular animals.

## RESULTS

### *Capsaspora* exhibits chemotaxis and chemokinesis in response to schistosome sporocysts

The initial report of *Capsaspora*, isolated from *B. glabrata* snails, revealed that *Capsaspora* migrated toward schistosome sporocysts. The sporocysts only attracted *Capsaspora* after they were initially attacked by *Capsaspora* and presumably began to ‘leak’ their contents into the media surrounding them.^15^ We repeated this analysis and indeed repeatedly observed migration of *Capsaspora* toward schistosomes that already had other *Capsaspora* cells attached (**Figure 1B, Videos S1–S6, Figure S1**).

Apart from the direction of movement, we asked if the velocity of *Capsaspora* movement was influenced by the presence of schistosome sporocysts. We discovered that *Capsaspora* showed increased overall cell movement (chemokinesis) when co-cultured with schistosome sporocysts (**Figure 1C**). Therefore, *Capsaspora* exhibits both chemotaxis and chemokinesis in response to leaking prey. Since chemokinesis is both less studied and easier to monitor than chemotaxis, we aimed to further investigate the chemokinesis behavior in this work.

### *Capsaspora* exhibits chemokinesis in response to cellular components from diverse organisms

We hypothesized that chemical components that leaked from the sporocyst were inducing *Capsaspora* chemokinesis. To test this hypothesis, we assessed if lysate from mechanically lysed prey could increase *Capsaspora* motility. We were unable to obtain enough schistosome sporocysts to test their lysate. However, *Capsaspora* has been shown to adhere and kill *Biomphalaria glabrata* embryonic (Bge) cells^15^—which are readily culturable in large quantities. Therefore, we tested if Bge cell lysate could induce *Capsaspora* chemokinesis. Indeed, we observed increased cell movement in the presence of Bge cell lysate in a dose-dependent manner (**Figure 1D, Figure S2A**). We further explored if other lysed cells induced chemokinesis. We tested lysates from both a mammalian cell line (HEK293T) and an insect cell line (JW18). Both induced chemokinesis **(Figure 1E, Figure S2B–C)**. These data together suggested that *Capsaspora* chemokinesis is induced by general cellular factor(s)—not a highly specific factor produced by only *Capsaspora*’s putative symbionts. To further probe the promiscuity of chemokinesis induction, we asked if even prokaryotic cell lysates could induce *Capsaspora* chemokinesis. We tested lysates of both a gram-negative bacterium (*Escherichia coli*) and a gram-positive bacterium (*Bacillus subtilis*). Both induced *Capsaspora* chemokinesis in a dose-dependent manner (**Figure 1F–G, Figure S2D–E**). In sum, *Capsaspora* increased its cell motility in response to soluble cell lysates from a remarkably wide range of cells across domains of life. Therefore, *Capsaspora* chemokinesis is induced by common and/or diverse cellular component(s).

### Bovine serum albumin is sufficient to induce chemokinesis in *Capsaspora*

Given that *Capsaspora* motility increases in response to such a range of lysates, we asked if it responded to an extracellular rich biological substance: fetal bovine serum (FBS). Remarkably, *Capsaspora* increased its movement in the presence of FBS in a dose-dependent manner, as well (**Figure 2A**). This observation further underscored the breadth of chemokinesis-inducing substances, and it afforded an abundant and stable chemokinesis-inducing material for further mechanistic studies.

**Figure 2.**
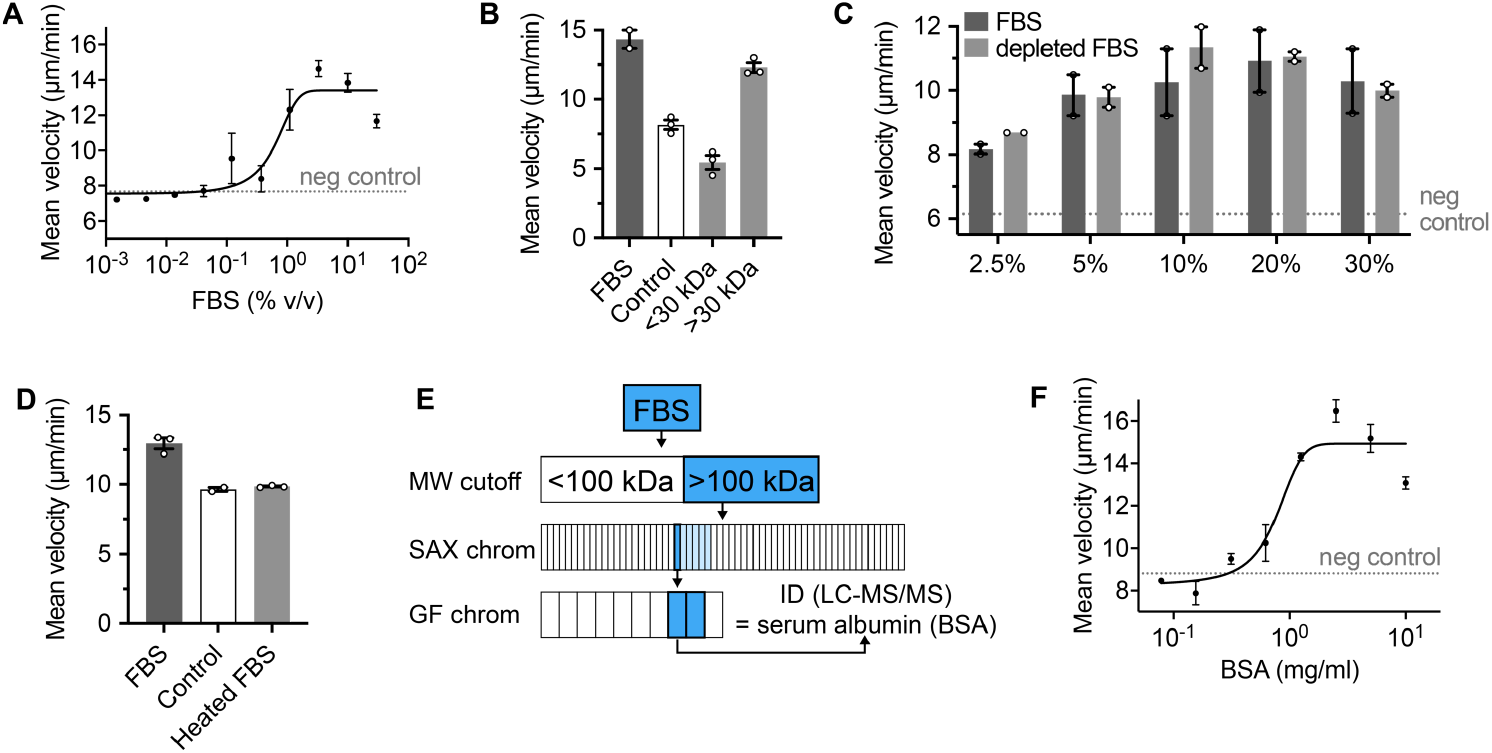
Bovine serum albumin is sufficient to induce *Capsaspora* chemokinesis. (A) *Capsaspora* motility upon addition of different concentrations of FBS. The final concentration of FBS in the assay is displayed on the x axis. Error bars represent standard error of the mean of a biological triplicate of three individual wells, each with dozens of cells (n=3). The dashed line indicates baseline motility upon addition of negative control (water). (B) *Capsaspora* motility upon addition of small molecule (< 30 kDa) or macromolecule (> 30 kDa) components of FBS, relative to full FBS and control (water). Error bars represent standard error of the mean of a biological triplicate of three individual wells of cells (n=3), except for the FBS positive control, which only had two wells. Individual biological replicates are displayed with white circles. (C) *Capsaspora* motility upon addition of different concentrations of FBS (dark grey) or lipoprotein-deficient FBS (light grey). Error bars represent the range of a biological duplicate of three individual wells, each with dozens of cells (n=2). Individual biological replicates are displayed with white circles. The dashed line indicates baseline motility upon addition of negative control (Chernin’s Balanced Salt Solution, CBSS^20^). (D) *Capsaspora* motility upon addition of heat-treated FBS, compared to untreated FBS and water (control). Error bars represent standard error of the mean of a biological triplicate of three individual wells (n=3), each containing dozens of cells. Individual biological replicates are displayed with white circles. (E) Bioassay-guided fractionation scheme to purify chemokinesis-inducing substance(s) from FBS. Boxes represent tested fractions from each separation. Dark blue boxes are the most active fractions, which were carried forward. Light blue boxes were less active fractions. Chromatograms, chemokinesis data for individual fractions, and SDS-PAGE image of final active fractions are reported in **Figure S3**. LC-MS/MS identification of major proteins is reported in **Dataset S1**. (F) *Capsaspora* motility upon addition of different concentrations of BSA. The final concentration of BSA in the assay is displayed on x axis. Error bars represent standard error of the mean of a biological triplicate of three individual wells, each with dozens of cells (n=3). The dashed line indicates baseline motility upon addition of negative control (water). Chemokinesis assay method A was used for panels A– C, E. Method B was used for panels D and F.

First, we aimed to identify a single pure component from FBS that was sufficient to induce chemokinesis. We investigated if the inducer was a small molecule or a macromolecule by separating FBS with a 30 kilodalton (kDa) molecular weight cutoff filter. We identified that only the > 30 kDa fraction induced chemokinesis—indicating that the inducer is macromolecular (**Figure 2B**). We previously found that serum lipoproteins trigger cellular aggregation in *Capsaspora*.^8,9^ To determine if these lipids were also important for chemokinesis, we tested lipoprotein-deficient FBS, which contains only ∼5% of the natural level of lipoproteins. We found that the depleted FBS induced chemokinesis with the same potency as full FBS, indicating that lipoproteins are not necessary for chemokinesis induction by FBS (**Figure 2C**). We further investigated if the active component was heat-labile. It was, suggesting the active component is a protein or heat-labile glycan (**Figure 2D**).

We fractionated the macromolecular material from FBS using a strong anion exchange (SAX) column (**Figure 2E, Figure S3**). We collected one of the most active fractions and subjected it to further fractionation by size exclusion column chromatography (SEC) (**Figure 2E, Figure S3**). Chemokinesis-inducing activity appeared to correlate with three proteins that migrated at ∼ 70 kDa, ∼60 kDa, and ∼40 kDa by sodium dodecyl sulfate-polyacrylamide gel electrophoresis (SDS-PAGE) (**Figure 2E, Figure S3I**). The primary proteins present in these gel bands were bovine serum albumin (BSA), alpha-fetoprotein (AFP), and alpha-2-HS-glycoprotein (fetuin A) (**Dataset S1**). These are all major components of fetal serum. BSA is readily available as a purified protein. Therefore, we investigated its ability to induce chemokinesis as a single pure component. We found that BSA could induce chemokinesis in a dose-dependent fashion (**Figure 2F**). Since FBS contains approximately 25 mg/ml of BSA,^19^ the EC_50_ value of FBS (∼1%, **Figure 2A**) should contain ∼0.25 mg of FBS—which is close to our observed EC_50_ of pure BSA (∼1 mg/ml, **Figure 2F**). Therefore, it is plausible that BSA is the primary (or sole) chemokinesis-inducing agent in FBS. It is also possible that AFP, fetuin A, and/or other proteins may also contribute in proportion to their concentration in FBS, as well.

### *Capsaspora* chemokinesis response is not specific to bovine serum albumin

Since diverse cell lysates (that do not contain BSA) increase *Capsaspora* motility, non-BSA components must also be chemokinesis inducers. Among other readily available proteins, human serum albumin and oval albumin also induced migration in *Capsaspora* in a dose-dependent fashion (**Figure 3A–B**). Oval albumin is especially notable, since it is not homologous to BSA. Gamma globulins also appeared to slightly induce motility at only a single concentration (**Figure 3C**). However, not all proteins increased *Capsaspora* motility—myoglobin failed to induce a significant increase in motility (**Figure 3D**).

**Figure 3.**
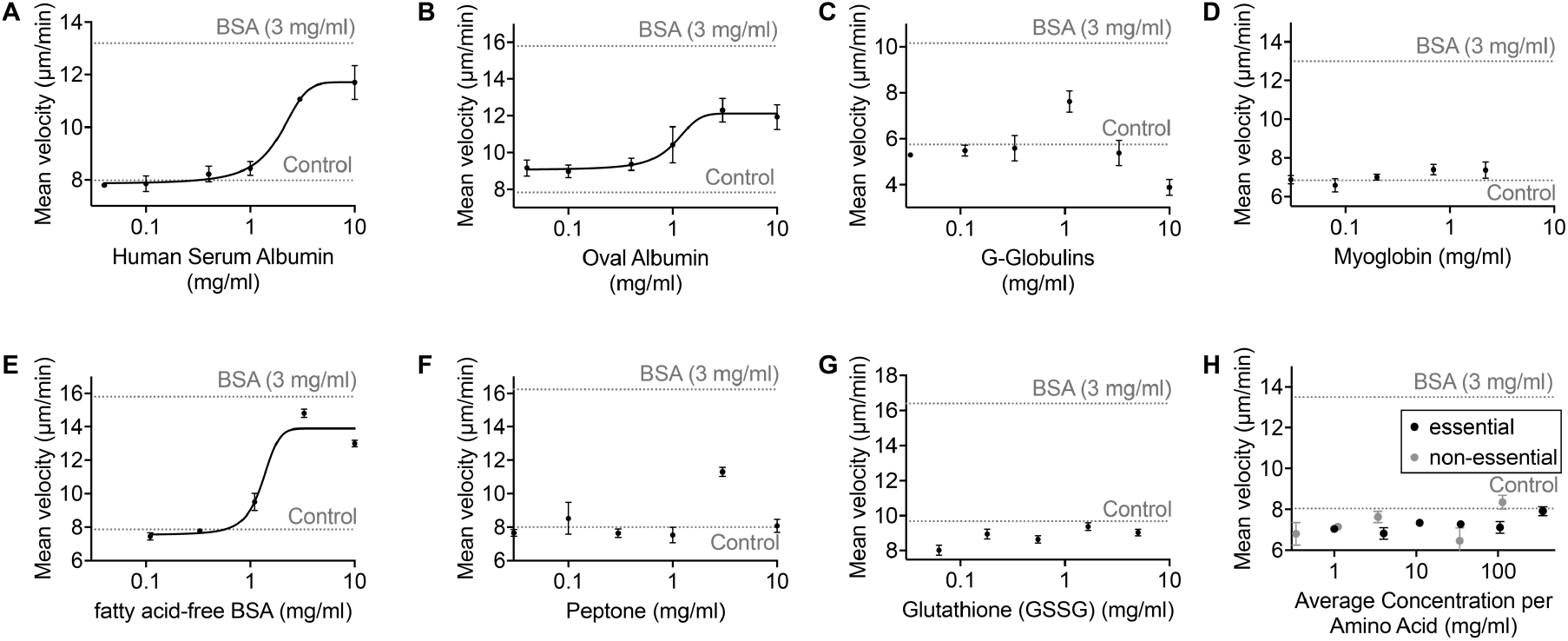
Other proteins increase *Capsaspora* motility. *Capsaspora* motility upon addition of different concentrations of (A) human serum albumin, (B) oval albumin, (C) G-globulins, (D) myoglobin, (E) fatty acid-free BSA, (F) peptone, (G) glutathione, and (H) mixtures of ‘essential amino acids’ and ‘non-essential amino acids’. In all cases, the final concentration of the inducer in the cell culture is displayed on x axis. Error bars represent standard error of the mean of a biological triplicate of three individual wells, each with dozens of cells (n=3). The dashed line indicates baseline motility upon addition of negative control (water). Chemokinesis assay method B was used for panels A–E, G. Method A was used for panels F and H.

We next considered which moieties of proteins were necessary to promote *Capsaspora* motility. First, the most active proteins are known to be carriers of lipids in the serum. To test if the protein-bound lipids were important for induction, we examined chemokinesis by fatty-acid free BSA.

This BSA induced motility with a similar potency as native BSA, suggesting that the fatty acids bound to BSA are not important for activity (**Figure 3E**). Next, we tested if whole protein was necessary or if individual peptides (or even amino acids) could induce chemokinesis. Often simple peptides are ligands for receptors that drive cellular responses. For example, human immune cells respond to N-terminal peptides of bacterial proteins, which contain *N*-formyl methionine.^21^ Therefore, we tested if individual peptone peptides could induce chemokinesis. They slightly induced motility at a single concentration (**Figure 3F**). However, another simple peptide (glutathione) failed to induce any aggregation (**Figure 3G**). Amino acids were incapable of inducing chemokinesis (**Figure 3H**). These data suggest that even some peptides may be sufficient to induce chemokinesis.

### Chemokinesis inducer does not substantially decrease adhesion of cells to surface

To investigate how these inducers increase *Capsaspora* motility, we first asked if BSA influenced the adhesion of *Capsaspora* to surfaces. Surface adhesion is intimately associated with motility. If cells stick too tightly to a surface, they cannot move.^22^ Alternatively, if they adhere too weakly, then mesenchymal-type motility is also impossible.^23^ Prior work has shown that extensive coating with BSA (for 1 hour) can decrease the adhesion of *Capsaspora* to untreated plastic surfaces.^14^ Therefore, we examined if BSA decreased *Capsaspora* adhesion in our chemokinesis experimental conditions (immediate analysis ∼2 minutes after addition of BSA to cells adhered to tissue culture-treated surfaces). We found no significant decrease in adherence upon addition of a chemokinesis-inducing concentration of BSA, followed by washing the cells with a level of force that removes approximately half of the surface-attached cells (**Figure 4A**). Although we cannot rule out a very slight inhibition of attachment speeding up *Capsaspora* movement, our data do not support this hypothesis.

**Figure 4.**
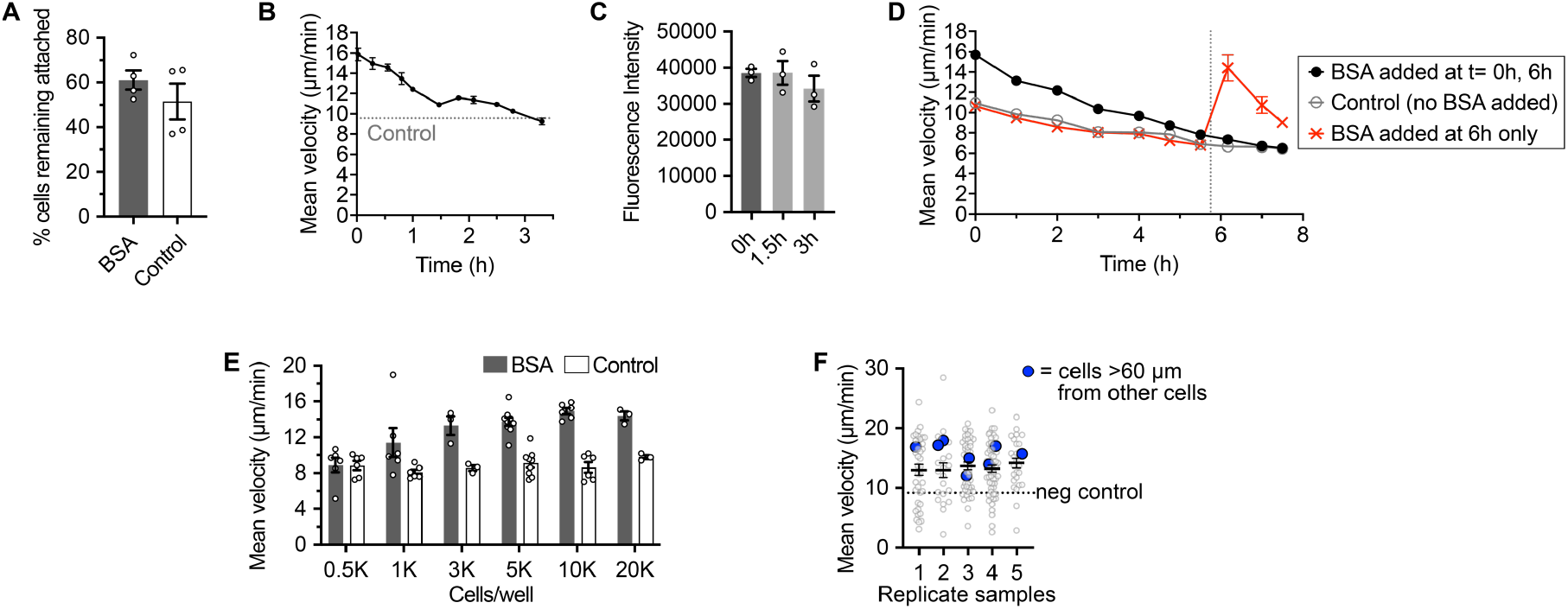
Insights into chemokinesis induction mechanism. (A) *Capsaspora* adhesion to tissue culture-treated plastic microplate wells upon addition of 3 mg/ml BSA or control (water). Error bars represent standard error of the mean of the means of four individual experiments (n=4). In each experiment, two or three wells were quantified for each condition. The means from each experiment are displayed as white circles. Data from the individual wells from each experiment are in **Figure S4**. (B) *Capsaspora* motility at different time points after addition of BSA (3 mg/ml). Error bars represent standard error of the mean of a biological triplicate of three individual wells, each with dozens of cells (n=3). The dashed line indicates baseline motility before addition of BSA. (C) Quantity of fluorescently labeled BSA remaining in the *Capsaspora* motility assay after 1.5 hours and 3 hours. Error bars represent standard error of the mean of a biological triplicate of three individual wells (n=3). Individual biological replicates are displayed with white circles. See raw gel image in **Figure S5B**. (D) *Capsaspora* motility at different time points after addition of BSA (3 mg/ml). Error bars represent standard error of the mean of a biological triplicate of three individual wells, each with dozens of cells (n=3). The black filled circles represent wells that received BSA (3 mg/ml) at the start and again after ∼6 hours. The open grey circles represent wells that received no BSA. The red “X” symbols represent wells that received no initial BSA treatment, but did receive BSA (3 mg/ml) after ∼6 hours. The dashed line indicates the time when the ∼6 hour BSA dose was added to the black and red conditions. (E) *Capsaspora* motility with different densities of cells upon addition of BSA (3 mg/ml) or control (water). Error bars represent standard error of the mean of two to six replicates individual wells, each with dozens of cells (n=3–9). Individual biological replicates are displayed with white circles. (F) Motility of individual *Capsaspora* cells from five replicate wells, highlighting cells that are distant from others (blue, incapable of cell-cell contact) exhibit increased motility on par with the average of their well. Circles represent each tracked cell, line and error bar represent mean and standard error of the mean for that well. Dashed line indicates the mean negative control migration in wells with water added instead of BSA. In all panels, the chemokinesis assays were performed using method B.

### Incubation with chemokinesis inducer desensitizes *Capsaspora* to further chemokinesis induction

We monitored the kinetics of the increased motility phenotype, and we found that it dissipated over the course of hours (**Figure 4B**). Initially we suspected that BSA might be depleted from the media, causing the decreased chemokinesis. To test this hypothesis, we monitored the depletion of fluorescently labeled BSA (fl-BSA) from the media. After confirming that the fl-BSA induces chemokinesis (**Figure S5A**), we collected aliquots of the cell culture media during the chemokinesis experiment conditions. We found that the fl-BSA was not significantly depleted on the timescale of decreased motility (**Figure 4C, Figure S5B**).

Alternatively, we hypothesized that *Capsaspora* becomes desensitized to its chemokinesis inducer over time. To test this hypothesis, we examined if re-addition of BSA would cause a renewed increase in motility. Even though this re-addition doubled the concentration of BSA, we found no increased motility relative to the untreated control (**Figure 4D**, black circles and grey open circles). This desensitization is almost certainly due to the initial BSA treatment, since cells that were not pre-treated with BSA were still capable of a large motility increase upon BSA addition at the later time point (**Figure 4D**, red “X” marks). In this case the rate of motility decreased more sharply over time—possibly due to the cells running out of nutrients. Ultimately, the chemokinesis induction was temporary and could not be renewed by re-addition of similar concentrations of BSA. This result may indicate an active adaptation of *Capsaspora* to the presence of the chemokinesis inducer, as many organisms adapt to the current concentration of a molecular cue.^17,18^

### Chemokinesis response is cell-density-dependent but not contact-dependent

Cell density influences the speed and migration behaviors of different cell types. For example, in multiple cancer cell lines, higher cell density causes faster cell migration compared to lower densities.^24,25^ In contrast, some cells in developing animals stop moving when they contact each other at high density.^26^ Therefore, we asked if *Capsaspora* cell density regulated its motility in response to BSA. We seeded cells at various cell densities and tested for chemokinesis response to BSA. All previous experiments in this manuscript employed cells at 5,000 to 10,000 cells per 100 μl volume in a microtiter well. Here we found that *Capsaspora* failed to increase motility at the lowest tested density (500 cells per 100 μl volume in a microtiter well, **Figure 4E**). Therefore, the chemokinesis response may depend not just on the presence of inducing environmental cues, but also on a *Capsaspora*-derived ‘signal’.

This ‘signal’ may be a soluble factor that is released from cells, or it may be mediated through contacts between adjacent cells.^27^ To assess the importance of cell-cell contacts, we examined if cells that were too distant from others for direct contact still exhibited chemokinesis. Prior work has shown that *Capsaspora* filopodia can be up to 24 μm long.^14^ We also have also not observed filopodia longer than this. Therefore, to conservatively bin cells that should not be contacting other cells, we monitored cells that were > 60 μm away from any other cell in the well. We compared the chemokinesis response of these ‘distant’ cells to the average of the cells in their well and found that the distant cells migrated as fast as the average (**Figure 4F**). Therefore, we conclude that the cell-derived chemokinesis signal is not dependent on cell-cell contacts. It may be a secreted soluble molecule.

In sum, it may be that the ultimate chemokinesis ‘signal’ is a soluble factor, which is produced by *Capsaspora* and secreted in response to the addition of the chemokinesis cue. Alternatively, the density-dependent signal may be constantly secreted by *Capsaspora* and synergize with the environmental cue to essentially provide an AND logic gate where both the *Capsaspora* signal and the environmental cue must be present for chemokinesis to proceed.

### Chemokinesis alone fails to significantly improve predation

Our experiments have found evidence that *Capsaspora* cells exhibit chemokinesis in response to various lysates and proteins. While chemotaxis can direct protist predators towards prey,^28^ the role of chemokinesis in predation is unclear. To explore this, we developed a mathematical model of *Capsaspora* cells responding to a lysed schistosome that exudes a chemokinesis-inducing chemical, see **Supporting Information 2**. The model was developed to explore the possibility that chemokinesis alone may prove beneficial for predation, in the absence of a chemotactic response.

Previous mathematical models have considered the effect of chemokinesis in concert with chemotaxis, e.g. with application to leukocytes^29^ and bacteria.^5,6,30^ The theoretical expectation is that organisms displaying chemokinesis in the absence of chemotaxis will accumulate away from the chemo-effector.^31^ High chemo-effector levels lead to increased locomotion speed, which enhances diffusion due to random cell motion. This should not be a useful response for predators, as it would cause them to diffuse more rapidly as they approach their prey, transporting them away from their target. However, *Capsaspora* may make use of chemokinesis in conjunction with irreversible attachment to aid predation. When *Capsaspora* cells reach the schistosome, they pierce the parasite and become irreversibly attached to it. Since chemokinesis does not alter a cell’s stochastic trajectory, but only the speed at which the latter is traversed, the number of cells that reach the schistosome are unchanged by speed of locomotion. However, coupled with irreversible attachment, the chemokinetically increased arrival rate at the schistosome implies a greater residence time for protists, which would be advantageous for predation.

To explore these hypotheses, we simulated a population of protists interacting with a schistosome, assumed spherical for simplicity. The schistosome was also assumed stationary, based on the very small movement of these organisms in our in vitro experiments. Further, the schistosome was assumed to exude chemicals with the same diffusivities, and eliciting the same chemokinetic responses for *Capsaspora*, as BSA or HSA. Details of the simulations can be found in **Supporting Information 2**. Results were obtained for simulated protists with and without a chemokinetic response. The simulations were restricted to a 15-minute period, to avoid the effect of cell clustering, which happens at longer times. As shown in **Figure 5A**, the simulations predict no significant difference between the distribution of residence times with and without chemokinesis. Thus, our model predicts that chemokinesis-inducing chemicals from a prey would not be directly beneficial to predation on their own.

**Figure 5.**
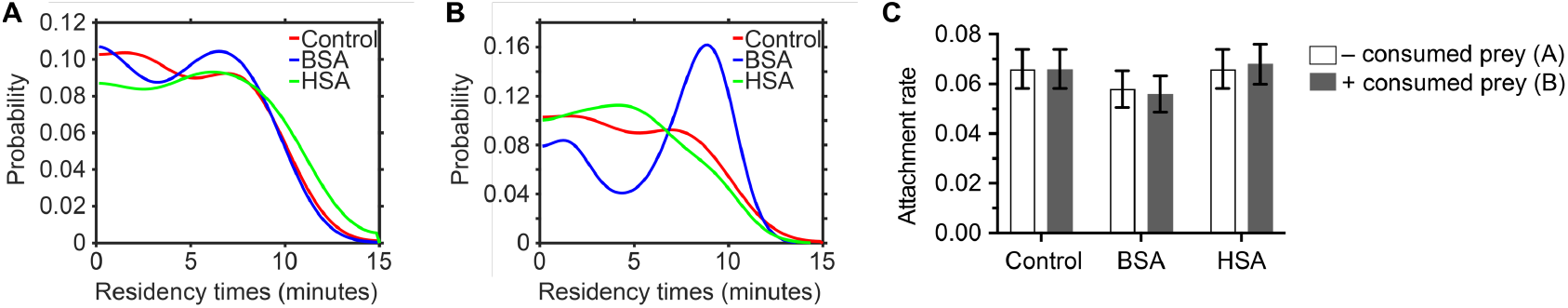
Numerical simulations of predator-prey interaction with chemokinesis. (A) Probability of different residency times over a 15-minute simulation period, from a kernel density estimate for a population of *Capsaspora* cells predating a schistosome with chemokinesis to BSA and HSA plus irreversible attachment. (B) is the same simulation as (A), but in the presence of recently consumed lysed schistosomes. (C) Attachment rates for protists predating a schistosome surface. Bars are errors propagated from the standard error of the mean (n=1000).

Another way that chemokinesis may aid predation is by increasing dispersion away from recently lysed prey. If the prey releases a large amount of chemokinesis-inducing chemicals, this would increase cell speed, and thus diffusion, encouraging higher dispersal away from the region where there is no longer prey. Thus, randomly moving protist cells need to search effectively less space before encountering another schistosome, with a benefit to predation through increased residency times. To explore this hypothesis, we repeated our simulations, this time using an initial chemical profile that models several recently lysed schistosomes in the vicinity of an intact one. The results of these can be seen in **Figure 5B**, which shows that, while HSA does not appear to have much effect on residency time, BSA increases the probability of higher residency times. The origin of this may lie in the different chemokinetic response curves for these two proteins (see **Figure 2F** and **3A**), with BSA having a significantly higher saturation speed than HSA. This means that similar concentrations induce a stronger increase in locomotion speed. In addition, we note that, since BSA and HSA diffuse much faster than the effective locomotion diffusivity of *Capsaspora*, the chemokinesis-inducing chemical will need to induce a strong response to aid predation before it rapidly diffuses into the surrounding medium.

Thus, our simulations indicate that, for sufficiently strong chemokinetic responses, chemokinesis may be beneficial to protist predators by increasing dispersal at sites of lysed prey, helping them to move more rapidly away from lysed prey and reach living prey faster. This may, however, not be sufficient to significantly impact predation. **Figure 5C** shows the attachment rate of protists in the presence and in the absence of recently-depleted schistosomes (which can enhance dispersion), for chemokinesis-inducing proteins and a chemokinesis-less control. In all cases simulated, only a maximum of just under 7% of the population reached the schistosome surface, with no significant difference between attachment rates with and without chemokinesis. Our model thus predicts that chemokinesis alone is likely not sufficient for successful predation. However, it may be that, acting in concert with chemotaxis, chemokinesis can provide an advantage over chemotaxis alone, as mathematical models have predicted in the case of bacteria.^5^ To show this, the chemoattractants to which *Capsaspora* responds would need to be identified and quantified, which is beyond the scope of this study. This would allow our mathematical model to be extended, parameterized, and compared with experimental results.

## DISCUSSION

Here, we have reported that *Capsaspora* exhibits chemokinesis in response to damaged schistosomes, to cell lysate from diverse organisms, and to bovine serum albumin (BSA). This discovery complements earlier work revealing *Capsaspora*’s ability to sense and respond to chemical factors from its snail host and schistosome prey. Namely, *Capsaspora* forms large cellular aggregates in response to lipids from its host snail,^8-10^ and it chemotaxes toward yet-unknown chemical cues from damaged schistosome prey (initially observed by Owczarzak et al.^15^ and quantified here in our work). Therefore, an expanding array of chemical factors from its host and prey shape the behavior of this microbial predator.

We were initially surprised that motility would increase in response to a wide diversity of protein cues from many sources. Even bacterial lysate was an inducer. *Capsaspora* has not yet been shown to prey on bacteria, but its filasterean relative *Ministeria vibrans* does.^32^ Responding to general cues can be advantageous to cells that promiscuously target diverse prey. For example, neutrophils have evolved to sense *N*-formylmethionyl peptides, which are universal in nascent bacterial proteins.^21^ Therefore, *Capsaspora*, which appears to promiscuously prey on many cell types,^15^ may benefit from sensing common and/or diverse prey cues.

We were also surprised by the relatively high concentration of chemokinesis inducer required for maximally increasing motility (∼3 mg/ml of BSA). Since the total protein content of cells is typically ∼200-300 mg/ml,^33^ nearby *Capsaspora* can certainly be exposed to prey-derived protein concentrations in excess of 3 mg/ml upon prey lysis. We speculate that this high concentration of protein could have influences beyond a classical ligand-receptor interaction with the *Capsaspora* cells. For example, the proteins may coat the surface of the substrate (or even surfaces of *Capsaspora* cells) to block ligand-receptor interactions, as well. Since surface adhesion is intimately related to motility rates, non-specific protein binding that improves or inhibits cell-substrate adhesion could play a role in the observed motility increases.^22,23,34^ We observed no significant influences on cell-substrate adhesion upon brief exposure to BSA; however, minor changes in adhesion that are below our limit of detection may be significant for motility rates. Further work is warranted to explore the role of cell-substrate adhesion in the increased motility of *Capsaspora* near prey. This work would ideally incorporate substrates that are relevant to the natural life of *Capsaspora* (beyond the tissue culture-treated surfaces and fibronectin-coated surfaces [**Figure S1B–C**] that we used in this work).

The observed cell-density dependence of the chemokinesis response is reminiscent of cell-cell communication behaviors during chemotaxis in neutrophils and *Dictyostelium discoideum*.^35,36^ In these cases, chemotaxing cells secrete their own signal molecules to relay the message to neighboring cells. In both cases, the ultimate outcome is a cooperative behavior. For neutrophils, this cooperation brings more immune cells to the site of an infection. For *D. discoideum*, a multicellular fruiting body is formed. Likewise, predation in *Capsaspora* is a cooperative behavior, where multiple cells can lyse larger prey and share the released nutrients more efficiently when they co-localize.^9,15^ Therefore, it is sensible for cell-cell signaling to contribute to *Capsaspora*’s regulation of predation. Ongoing work is aimed at characterizing the identity of the putative secreted signal and comparing the response mechanism to that in cell-cell communication in extant animals.

Finally, chemokinesis is a mysterious phenotype. In contrast to chemotaxis, it does not obviously lead to improved predation—although many predators and symbionts do employ it.^6,30,37,38^ On its own, a gradient of a chemokinesis-inducing agent leaking from prey may actually be detrimental to predation—causing accumulation of cells in regions further away that induce lower motility.^34^ In our model of *Capsaspora* motility, we found that chemokinesis alone was unable to significantly improve *Capsaspora* predation. We considered the possibility that the long-term attachment of *Capsaspora* to prey may be sufficient to make chemokinesis a beneficial trait; however, we found that to be untrue. It appears likely that chemokinesis is only a beneficial predatory trait when it synergizes with chemotaxis.^5^ Thus the ability of *Capsaspora* to chemotax may establish the benefit of chemokinesis. Given our demonstration of chemokinetic responses of *Capsaspora* to bacterial lysates, in future studies it would be interesting to apply our simulation framework to explore the role of chemokinesis in the predation of bacteria by protists that do not obviously make use of chemotaxis, such as *Acanthamoeba*.^39^

## CONCLUSION

In sum, we have found that a predatory microbe’s motility is increased by proteins released from leaking prey. The behavior is density-dependent, suggesting cooperative predatory behavior. This increased motility near prey is not expected to benefit the predator on its own. However, since chemotaxis also occurs, the combined effect of chemotaxis and chemokinesis is likely beneficial for predation.

## METHODS

### Cell strain and growth conditions

*Capsaspora owczarzaki* cell cultures (strain American Type Culture Collection ATCC®30864) were grown axenically in 25-cm^2^ culture flasks with 6 ml ATCC media 1034 (modified PYNFH medium) containing 10% v/v heat-inactivated Fetal Bovine Serum (FBS, Corning 35-011-CV), hereafter growth media, in a 23 °C incubator. Cells were obtained during exponential growth phase by passaging ∼0.9 – 1.2 ml of adherent cells scraped into suspension within the specified density range, into 5ml of growth media. The culture was incubated at 23 °C for 24 h to achieve desired density range of 3 × 10^6^ to 20 × 10^6^ cells/ml.

### *S. mansoni* miracidia isolation and transformation into primary sporocysts

*S. mansoni* miracidia were isolated according to published protocols.^40,41^ Briefly, 3 infected mice livers were obtained from the Biomedical Research Institute (Rockville, MD) (BRI) Schistosomiasis Resource Center. They were shipped overnight with cold packs and added to ice-cold 1.2% sterile NaCl solution immediately after arrival using a sterile 50 ml centrifuge tube. The tubes with the livers were shaken vigorously, and any floating debris and the saline solution was aspirated. The livers were then poured into the ice-cold cup of a Waring blender with another 30 ml of ice-cold sterile 1.2% NaCl solution and blended for 15–25 seconds non-stop making sure the livers were well-blended. The blended livers were centrifuged at 800 × g for 10 min at 4 °C. The supernatant was discarded, and the pellets were washed with ice-cold sterile 1.25% NaCl solution. The pellets were then resuspended with 40 ml room temperature sterile artificial pond water (recipe from BRI, 0.46 μM FeCl_3_, 220 μM CaCl_2_, 100 μM MgSO_4_, 310 μM KH_2_PO_4_, 14 μM (NH_4_)_2_SO_4_ in water adjusted to pH 7.2 with NaOH) and transferred into the aluminum-covered sterile volumetric flask. The flask was carefully filled with sterile pond water to approximately 3 cm above the foil covering the neck. The flask was covered with a small petri dish and a bright light shone across the top of the flask. The eggs started to hatch at room temperature. When miracidia had amassed at the surface after 10 min, around 10 ml of the pondwater with miracidia was transferred to a sterile petri dish. The collected miracidia were centrifuged at 290 × g at 4 °C for 2 minutes. Only 1 ml of pondwater was left with the miracidia pellet and another 1ml of Chernin’s Balanced Salt Solution (CBSS+)^20^ was added to the petri dish. The miracidia were then transferred to a 24-well plate and incubated at 26 °C overnight to stimulate miracidial transformation into primary sporocysts.

### Chemotaxis/chemokinesis near schistosomes

*Capsaspora* cells were counted and resuspended such that the final cell density was 60 × 10^3^ cells/ml. 200 μl of the *Capsaspora* cells were seeded in each well of a TC-treated 96-well plate (Corning #353219) and incubated overnight at 26 °C. In the experiment run on fibronectin-coated plated (Fig S1B–C), a TC-treated 96-well plate (Corning #353219) was pre-coated with 100 μl of 70 μg/ml fibronectin (Sigma #F1141) in 25% FBS-free media before adding *Capsaspora*. The next day, schistosome sporocysts were centrifuged for 30 seconds at 300 × g at room temperature and the supernatant was aspirated immediately. The pellet was then resuspended with 650 μl 25% FBS-free media. Media was aspirated out of the *Capsaspora* wells and replaced with 200 μl of the sporocyst cell culture to afford a few schistosomes per well. Negative control wells received media without schistosomes. For imaging each well, a field of view was set such that there is at least one sporocyst surrounded by many *Capsaspora* cells. Images were collected every one minute for 4 hours using an A1 Nikon inverted microscope. Cells were then analyzed using Imaris image analysis software (Bitplane, version 10.0.1). ImageProcess was used to invert the image, the schistosome was set as the origin reference, and spots were created. Estimated cell diameter was set to 6.43 μm. Background was subtracted, and the automatic threshold was used to quality filter the spots. Cells 70–240 μm from the origin reference frame were selected. The autoregressive motion algorithm was used for tracking. To remove spurious readings by floating cells (which appear to migrate abnormally quickly and/or appear/disappear quickly), we excluded cells that migrated > 11.2 μm in adjacent frames or lost tracking for > 3 frames. The ‘fill gap’ function was disabled, and only tracks ≥3 frames were analyzed. For chemotaxis, the distance moved relative to the reference (schistosome) was reported for each cell track. For chemokinesis, the distance moved frame-to-frame for each cell in each frame was reported.

### Chemokinesis assay

This assay was always performed with the strain ATCC®30864 unless mentioned otherwise. The standard chemokinesis assay was performed on TC-treated 96-well plate (Corning #CLS3997). Either 5 × 10^3^ cells or 10 × 10^3^ cells were seeded in 90 μl of FBS-free growth media per well in the TC-treated 96-well plate and allowed to settle for 2 h.

### Method A

After seeding cells and settling for 2 h, 85–90% of the FBS-free media was removed by aspiration and replaced with 85 μl Chernin’s balanced salt solution (CBSS+). Samples were then added at 10 μl volume such that the total volume in each well was 100 μl. At concentrations tested higher than 10%, less CBSS+ was added, and a larger sample volume was added to reach the desired concentration with a final volume of 100 μl. Chemokinesis activity was observed right after the sample addition by taking bright field images every 5 seconds for 3 minutes using an Agilent BioTek Cytation 10 confocal imaging reader.

### Method B

After seeding cells and settling for 2 h, samples were directly added at 10 μl volume such that the total volume in a well is 100 μl. Chemokinesis activity was observed right after the sample addition by taking bright field images every 5 seconds for 3 minutes using an Agilent BioTek Cytation 10 confocal imaging reader.

### Chemokinesis quantification

When the Cytation 10 was used to record cell movement, analysis was performed using BioTek Gen 5 software. Briefly, images were pre-processed using a dark background with background flattening using a rolling ball diameter of 9 μm and image smoothing strength of 2 cycles. Kinetic frame alignment was employed based on the previous valid image with a maximum offset of 200 μm. The cellular analysis module was employed to track cells. A primary mask was applied using a threshold value of 5000 with a dark background, split touching objects, track objects, and fill holes in masks. Object size was filtered to be 1–100 μm. To remove spurious readings by floating cells (which appear to migrate abnormally quickly and/or appear/disappear quickly), we made a subpopulation of cells that migrated ≤60 μm and were tracked for ≥ 18 frames. The ‘average velocity’ of this subpopulation of cells was reported as the ‘mean velocity’.

### Cell lysis

*Biomphalaria glabrata* embryonic (Bge) cells^42,43^ were grown in Bge medium with 10% FBS at 26 °C. The culture was resuspended to 20 × 10^3^ cells/ml and snap-frozen in 20 μl aliquots until further use. The frozen aliquot was freeze-thawed 5 times with a 20 second thawing period between each cycle. An additional 80 μl of FBS-free media was added to make the final volume 100 μl. The lysed culture was centrifuged at 15,000 × g for 5 minutes and the supernatant was collected for further testing.

Both animal cell cultures were freshly received and resuspended to appropriate densities using FBS-free media. The cultures were then centrifuged at 15,000 × g for 5 min and the cell pellets were lysed using freeze-thaw method for 5 times with 20 second thawing period between each cycle. The lysed cultures were centrifuged at 15,000 × g for 5 min to collect the lysate supernatant.

*E. coli* MG1655 culture was grown overnight in lysogeny broth (LB) at 37 °C with shaking and sub-cultured to 50 ml with 1:50 dilution of the overnight culture into fresh media. The 50 ml culture was incubated an additional 18 hours with shaking at 37 °C. The culture was centrifuged at 10,000 × g for 15 min. The cell pellet was resuspended in 1 ml of FBS-free media. The culture was lysed using a probe sonication for 10 seconds followed by a 20-second paused, repeated for a total of 5 min. The lysed culture was centrifuged at 10,000 × g for another 15 min and the lysate supernatant was collected for further analysis.

*B. subtilis* 168 culture was grown overnight in lysogeny broth (LB) at 37 °C with shaking and sub-cultured to 50 ml with 1:50 dilution of the overnight culture into fresh media. The 50 ml culture was incubated an additional 18 hours with shaking at 37 °C. The 50 ml culture was centrifuged at 10,000 × g for 15 min. The cell pellet was resuspended in 1 ml of 4 mg/ml lysozyme and incubated for 15 min. The culture was centrifuged again at 15,000 × g for 15 min and the cell pellet was resuspended in 1 ml of FBS-free media. The culture was lysed using sonication for 10 seconds followed by a 20-second paused, repeated for a total of 5 min. The lysed culture was centrifuged at 10,000 × g for another 15 min and the lysate supernatant was collected for further analysis.

### FBS Heat-treatment

100 μl of FBS in an Eppendorf tube was heated on dry heat block at 75 °C for 30 minutes. The treated and non-treated FBS was tested for chemokinesis assay. The FBS-free growth media was used as negative control.

### Bioassay-guided fractionation

FBS was initially fractionated using either Amicon Ultra 30 kDa or 100 kDa cutoff filters (Sigma, #UFC5030, #UFC5100) according to the manufacturer’s direction. Briefly, the filters were first washed with 500 μl of CBSS by centrifugation at 14,000 × g for 15 min. Then 500 μl of FBS was added to the cutoff filter and centrifuged at 14,000 × g for another 15 min. The small fractions (< 30 kDa, < 100 kDa) were collected for testing, and 20 μl of each large fraction (> 30 kDa or > 100 kDa) was added to 180 μl of CBSS for testing. 10 μl of each stored fraction was used for testing chemokinesis.

### Isolation using strong anion exchange column (SAX)

#### Chromatography sample preparation

20 ml of FBS was fractionated using an Amicon Ultra 100 kDa cutoff filter (Sigma, #UFC8100) according to the manufacturer’s direction. Briefly, the filters were first washed with 5 ml CBSS+ by centrifuging at 4,000 × g for 25 min. Then 10 ml of FBS was added to each cutoff filter and centrifuged at 4,000 × g for 50 min. The > 100 kDa was washed by adding 5 ml of buffer A (50 mM Tris Ethanolamine, pH 9.5), centrifuged at 4,000 × g for 25 min, and the concentrated sample was collected. The > 100 kDa fraction was then sterile-filtered using Costar Spin-X Centrifuge filters (Corning, #29442-752).

#### Chromatography Method

A GE HealthCare HiPrep^TM^ Q XL 16/10 column (Fisher, #45-002-068) was used. For the mobile phase, 50 mM Tris Ethanolamine, pH 9.5 was used as buffer A and 50 mM Tris Ethanolamine supplemented with 1 M NaCl, pH 9.5 was used as buffer B with a flowrate of 5 ml/min. The column was first washed with buffer B for 5 min and equilibrated with buffer A using 5 column volumes (CV). The 5 ml sample loop was then washed with buffer A followed by injecting 1.2 ml of the sample (> 100 kDa). Unbound material was washed from the column using 10 CV buffer A. The fractions were then eluted by using a linear gradient from 0– 55% B using 30 CV and 55–100% B using 20 CV followed by holding 100% B for 10 CV. All fractions including the flow through were collected using a fixed volume of 5 ml per fraction.

#### Chemokinesis sample preparation

The unbound fraction was pooled and labelled as fraction 1. To simplify, a systematic pooling strategy was employed. Initially, a primary pool was made by combining 50 μl from each of the 30 consecutive eluted fractions and labelled them sequentially from fraction 2 through fraction 9. Each of these pooled fractions were concentrated using the 100 kDa cut off filters, centrifuged at 14,000 × g for 15 min at 10 °C. > 100 kDa fractions were then washed by adding 300 μl of CBSS and centrifuged again at 14,000 × g for 15 min at 10 °C and collected the > 100 kDa fractions. 20 μl of each of these fractions were added to another 70 μl of CBSS to such that the total volume was 90 μl and stored at 4 °C until testing.

Once the active pooled fractions were identified (pools 3 and 4), secondary pools were made by combining 100 μl from 10 consecutive initial fractions and labeled them sequentially. For example, fraction 31 contained original fractions 31–40, while fraction 33 contained fractions from 51–60. The same approach was applied to the other active initial pool 4. Pool 4 was subdivided into fractions 41, 42, and 43, where fraction 42 contained original fractions 71–80. The combined fractions were concentrated and washed as above. After collecting the > 100 kDa fractions, 20 μl of each of these fractions were added to 180 μl of CBSS to such that the total volume was 200 μl. These were stored at 4 °C until testing.

Upon identifying activity in specific secondary pools (pools 33 and 41), a tertiary pooling step was performed. Each active secondary pool was further divided by combining 50 μl from 5 consecutive initial fractions into a new pool and labeled accordingly. For example, fraction 33 was divided into 331 and 332, where fraction 332 contained original fractions 56–60 and fraction 421 contained original fractions 71–75. This iterative process allowed for the stepwise narrowing of active fractions. 20 μl of each of these fractions were added to another 80 μl of CBSS such that the total volume was 100 μl. These were stored at 4 °C until testing.

10 μl from each of these pools were added to the chemokinesis assay and CBSS was used as negative control

### Isolation using gel filtration (size exclusion chromatography [SEC])

#### Chromatography sample preparation

2 ml of each fraction contributing to the 331 pool (i.e. 51–55) was combined to achieve a final volume of 10 ml to prepare for SEC. First, the 100 kDa cutoff filter was washed with 5 ml CBSS+ by centrifuging at 4,000 × g for 25 min. The sample was then centrifuged at 4,000 × g for 35 min using 100 kDa cutoff filter. To the > 100 kDa fraction, another 3 ml of CBSS was added to the sample and centrifuged for another 30 min using 100 kDa cutoff filter and collected the final > 100 kDa for SEC fractionation.

#### Chromatography Method

A HiLoad 16/600 Superdex 200 pg (GE healthcare #28989335) column was used. For the mobile phase, 50 mM Tris Ethanolamine supplemented with 1 M NaCl, pH 9.5 was used with a flowrate of 1 ml/min. The column was first washed with the buffer for 5 min and equilibrated with the buffer using 0.2 column volumes (CV). The 1 ml sample loop was then washed with the buffer followed by injecting 1 ml of the sample (> 100 kDa). The fractions were then eluted by using isocratic condition using 5 CV. All fractions were collected using a fixed volume of 3 ml per fraction.

#### Chemokinesis sample preparation

500 μl from each fraction containing proteins (5–12) was individually centrifuged at 4,000 × g for 15 min at 10 °C using 100 kDa cutoff filters. The >100 kDa fraction was washed once with 200 μl CBSS and the remained 20 μl of >100 kDa fraction was directly tested for chemokinesis assay and stored at 4 °C for further analysis.

### Different cell density seeding

After measuring the cell density in a culture flask, the culture was appropriately diluted such that seeding with 90 μl of each diluted culture achieves the desired density.

### Desensitization experiments

Chemokinesis assay was performed using the method B and Gen5 for imaging analysis. 10 μl of 30 mg/ml BSA was added to induce chemokinesis as mentioned at t =0 and 6h.

### Adhesion assay

5 × 10^3^ cells were seeded in 90 μl of FBS-free growth media per well in the TC-treated 96-well plate and allowed to settle for 2 h. After seeding cells, 85–90% of FBS-free media was removed by aspiration and replaced with Chernin’s balanced salt solution (CBSS+). A Cytation 10 imaging reader was set to capture ∼80% of the well’s area to count this initial cell number. After imaging, 10 μl of either BSA or water were added to each well and allowed to sit for 2 min (to mimic the chemokinesis conditions). 85–90% of FBS-free media was removed by aspiration and replaced with 85 μl CBSS+ solution. The Cytation 10 imaging reader was again used to capture ∼80% of the well’s area to count the remaining cell number. The remaining cell number was divided by the initial cell number to determine the % cells remaining for each well.

### Numerical modelling of predator prey interaction with chemokinesis

Simulation of a population of *Capsaspora* cells was completed in MATLAB; details can be found in **Supporting Information 2**. *Capsaspora* cells were modelled as individual agents, using Monte-Carlo methods to account for stochasticity. Rules of motion were inferred using a data driven approach. We analyzed the motility statistics of WT *Capsaspora* cells on a uniform substrate from Video 1 in reference.^14^ We found that *Capsaspora* cells display strong directional persistence following a truncated normal distribution for the correlation between direction of currently extended filopodia (**Figure SII**.**3**) and the direction of the next extended filopodia. This implies that cells tend to carry on travelling in the same direction with only minor adjustments to the right or left, making larger adjustments rarely, so they would take a long time to reverse direction. We then measured distance travelled after extension of filopodia and found it follows an exponential distribution (**Figure SII**.**2**), so that distances travelled following filopodia extensions are fairly uniform. To model chemokinesis, we parameterized the curves found for BSA as well as HSA, which were found to have the same shape with the only difference being BSA saturates at a higher locomotion speed, details can be found in **Supporting Information 2**.

The schistosome was then modelled as a stationary circle, that would cause cells to become irreversibly attached if they made their way to the schistosome surface. The schistosome would then exude the chemokinesis-inducing chemical at a rate dependent on the local attachment rate of *Capsaspora* cells to the surface. Dispersion of chemical was modelled in a continuum as standard isotropic diffusion with a constant diffusivity. These simulations were run with and without chemokinesis to determine if chemokinesis was beneficial to predation by increasing residency time of *Capsaspora* at the schistosome surface. The simulations were then run again for the case of a chemical profile reflecting several recently lysed schistosomes, to determine if they would act as a ‘dispersal agent’ and encourage cells to find new prey.

### Reagents used in this study

| Reagent | Cat. No. # | Company |
| --- | --- | --- |
| TC-treated 96-well plate | CLS3997 | Corning |
| TC-treated 96-well plate | 353219 | Corning |
| FBS | 35-011-CV | Corning |
| Lipoprotein-deficient FBS | S5394 | Sigma |
| Ethanolamine | E9508-1L | Sigma |
| Bovine serum albumin | A9418-5G | Sigma |
| Human serum albumin | A1653-1G | Sigma |
| Oval albumin | A5503-1G | Sigma |
| G-globulins | G7516-1G | Sigma |
| Myoglobin | M0630-250MG | Sigma |
| Fatty acid-free BSA | 126609-5G | Sigma |
| Poly-L-Lysine | P4832 | Sigma |
| Glutathione | AAJ6216614 | Fisher |
| Peptone | DF0118-17-0 | Fisher |
| Essential amino acid mix | 11-130-051 | Fisher |
| Non-essential amino acid mix | 11140050 | ThermoFisher |
| Fluorescent BSA | A9771-100MG | Sigma |
| Fibronectin | F1141 | Sigma |

## SUPPORTING INFORMATION

Supplemental Appendix includes Supplemental Figures S1–S5.

Supplemental Information 2 includes details of the motility model.

Supplemental Videos S1–S6 are available on figshare (doi: 10.6084/m9.figshare.28847960).

Supplemental Dataset S1.

Raw image stacks for all chemokinesis assays and chemokinesis model code are available upon request.

## ACKNOWLEDGEMENTS

We thank the Schistosomiasis Resource Center for provision of schistosomes and Bge cells, which were provided by the Schistosomiasis Resource Center of the Biomedical Research Institute (Rockville, MD) through NIH-NIAID Contract HHSN272201700014I NIH: We thank Margaret Mentink-Kane and André Miller for instruction on schistosome isolation. We thank the Light Microscopy Center at Indiana University for support in image acquisition and analysis (funding provided by the NIH grant NIH1S10OD024988-01). We also thank the Indiana University Laboratory for Biological Mass Spectrometry and Jon Trinidad for proteomics assistance. We thank Emily Layton and Richard Hardy for providing mammalian and insect cell pellets. We also thank the entire Gerdt lab for insights and support that helped advance this project. This work was supported by the National Institutes of Health (R35GM138376) to J.P.G. The content of this paper is solely the responsibility of the authors and does not necessarily represent the official views of the National Institutes of Health. This work was also supported by a Camille Dreyfus Teacher-Scholar Award (TC-24-028), the Engineering and Physical Sciences Research Council [EP/W524700/1] and the Human Frontier Science Program (No. LT000941/2021-C).

