## Supplemental Appendix for "Chemokinesis by a microbial predator"

**Contents:** Supplemental Figures S1–S5.

### SUPPLEMENTAL FIGURES

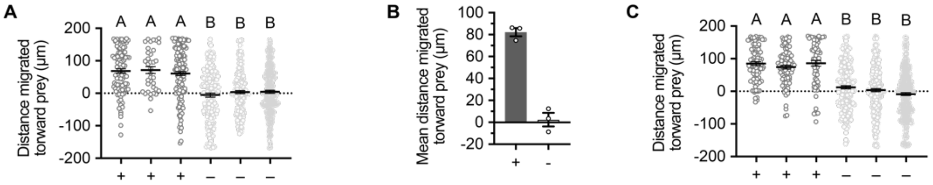


**Figure S1.** ***Capsaspora*** **chemotaxis to schistosome prey.** (A) Migration of individual cells toward schistosome prey (same data that is summarized in **Figure 1B**). The three left samples have a schistosome present. The three right samples do not ('prey' location was a randomly chosen spot in the middle of the field of view). Lines indicate means, and error bars represent standard error of the mean. Each circle is a single tracked cell. One-way ANOVA with multiple comparisons was performed, showing that all samples with a schistosome had significantly higher motility toward the schistosome than all three samples lacking a true schistosome. (B–C) Repeated chemotaxis experiment using fibronectin-coated dishes instead of plain tissue culture-treated surfaces (parallel to **Figures 1B and S1A**). (B) Chemotaxis of *Capsaspora* toward schistosome sporocysts. Average net movement toward the schistosome is reported (negative value would indicate movement away from schistosome). Error bars represent standard error of the mean of a biological triplicate of three individual wells of cells (n=3). Individual biological replicates are displayed with white circles. Within each biological replicate, many individual cells were tracked. (C) Migration of individual cells toward schistosome prey (same data that is summarized in **Figure S1B**). The three left samples have a schistosome present. The three right samples do not ('prey' location was a randomly chosen spot in the middle of the field of view). Lines indicate means, and error bars represent standard error of the mean. Each circle is a single tracked cell. One-way ANOVA with multiple comparisons was performed, showing that all samples with a schistosome had significantly higher motility toward the schistosome than all three samples lacking a true schistosome.

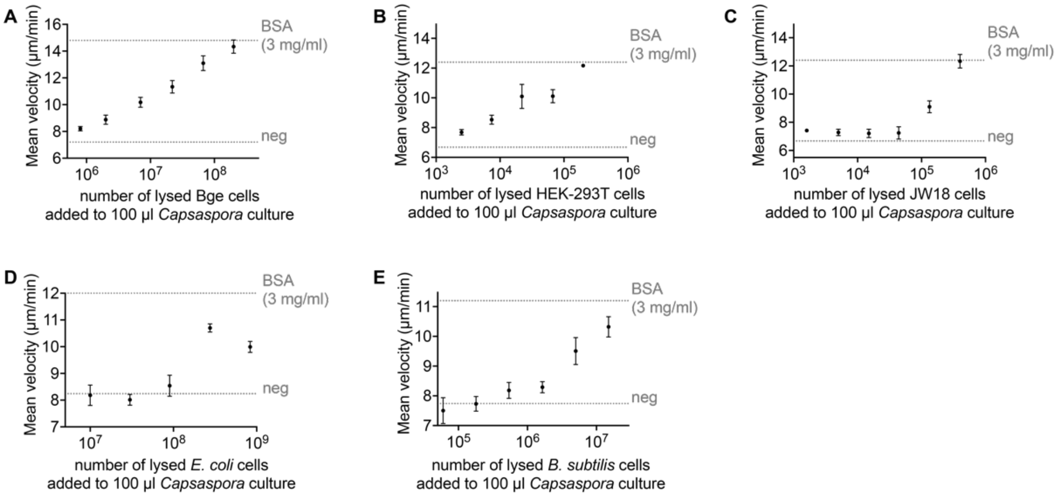


**Figure S2. *Capsaspora* chemokinesis is induced by cell lysates in a dose-dependent manner.** Data match those plotted in **Figure 1C–G**, where the data from only the most active concentration was plotted for each lysate. In all cases except for *E. coli* lysate, this was the highest concentration tested. (A) *Capsaspora* motility upon addition of different concentrations of Bge cell lysate. (B) *Capsaspora* motility upon addition of different concentrations of HEK-293T cell lysate. (C) *Capsaspora* motility upon addition of different concentrations of JW18 cell lysate. (D) *Capsaspora* motility upon addition of different concentrations of *E. coli* cell lysate. (E) *Capsaspora* motility upon addition of different concentrations of *B. subtilis* cell lysate. In all cases, the final concentration of lysed cell material added in the cell culture is displayed on the x-axis. Error bars represent standard error of the mean of a biological triplicate of three individual wells, each with dozens of cells (n=3). The dashed lines indicate baseline motility upon addition of negative control (water) or induced motility upon addition of 3 mg/ml BSA.

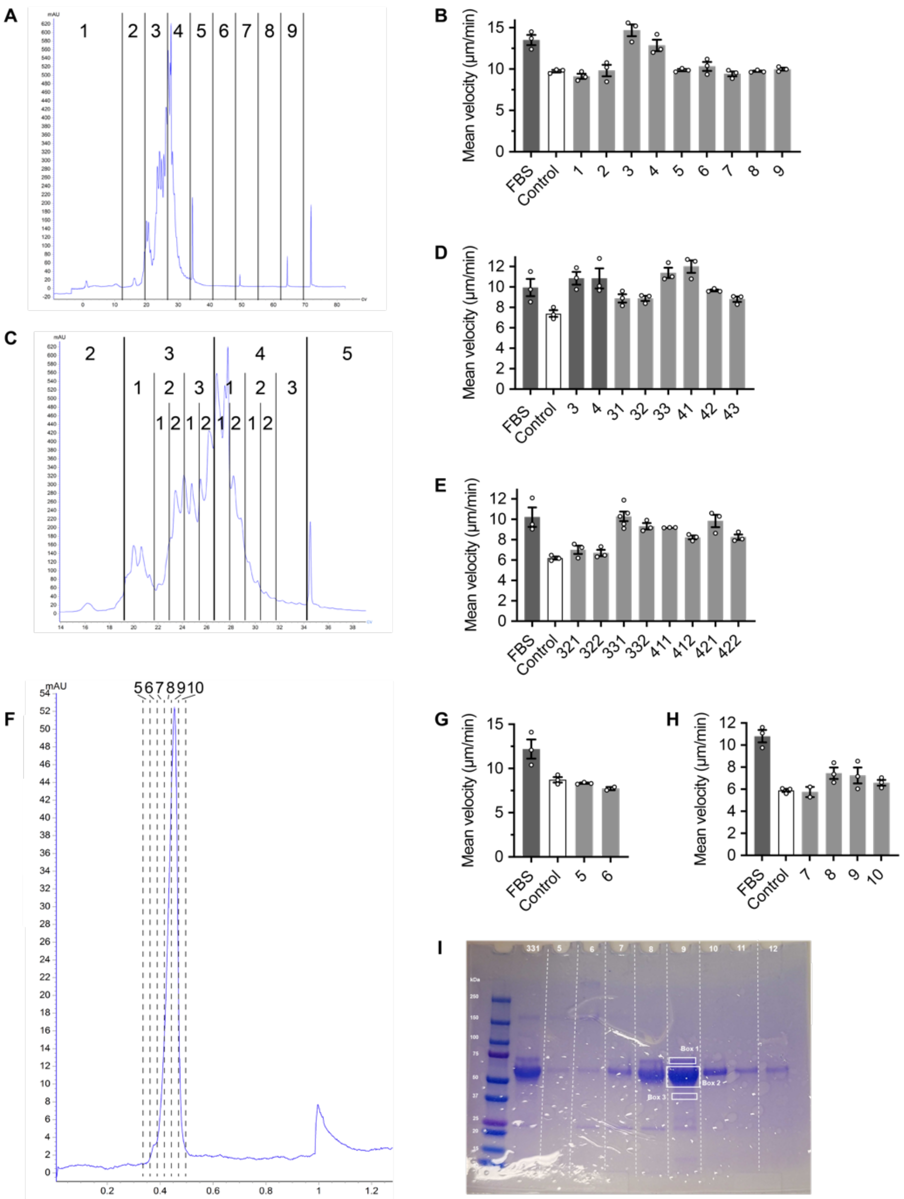


**Figure S3. Bioassay-guided fractionation of FBS.** (A) Chromatogram of FBS separated by anion exchange (AEX) chromatography. Coarse pooled fractions are shown. (B) *Capsaspora* motility upon addition of coarse fractions from AEX. (C) Chromatogram of FBS separated by anion exchange (AEX) chromatography. Medium-sized and fine-sized pooled fractions are shown. (D) *Capsaspora* motility upon addition of medium-sized pooled fractions from AEX. (E) *Capsaspora* motility upon addition of fine-sized pooled fractions from AEX. (F) Chromatogram of FBS separated by size exclusion chromatography (SEC). Fractions collected are shown. (G) *Capsaspora* motility upon addition of fractions 5 and 6 from SEC. (H) *Capsaspora* motility upon addition of fractions 7–10 from SEC. (I) Coomassie-stained SDS-PAGE gel of SEC fractions. Boxes are placed around gel bands that were excised for identification by tryptic peptide LC-MS/MS. For all motility assays, positive control (10 µl whole FBS) and negative control (10 µl CBSS) were included on the same day as the co-plotted sampled. In all plots, error bars represent standard error of the mean of a biological triplicate (in a couple cases duplicate) of individual wells, each with dozens of cells (n=3). Individual replicates are displayed with white circles.

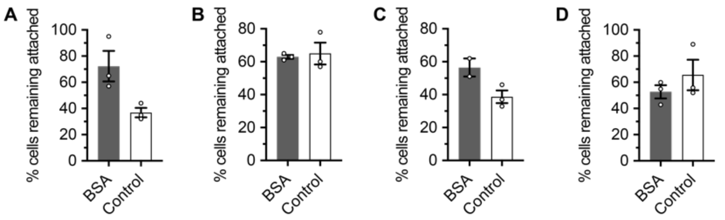


**Figure S4. Replicate experiments testing impact of BSA on *Capsaspora* surface adhesion.** *Capsaspora* adhesion to tissue culture-treated plastic microplate wells upon addition of 3 mg/ml FBS. Each panel is an experiment run on a separate day. The mean from each experiment was used to generate **Figure 4A**. Error bars represent standard error of the mean of two or three individual wells, each containing dozens of cells. Individual biological replicates are displayed with white circles.

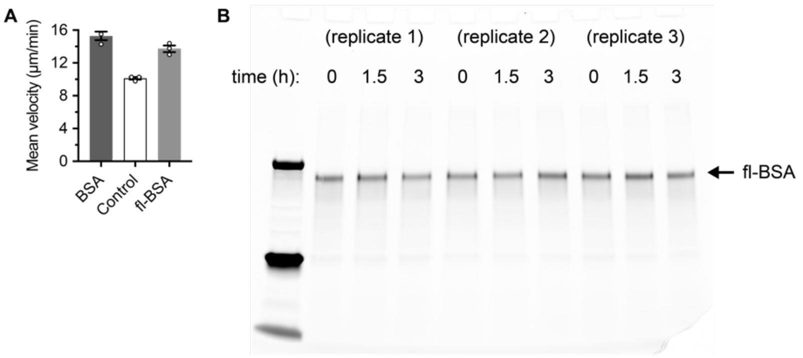


**Figure S5. Chemokinesis induction and lack of depletion of fluorescent BSA.** (A) *Capsaspora* motility upon addition of 3 mg/ml of BSA or fluorescently modified BSA (fl-BSA), compared to a negative control (water). Error bars represent standard error of the mean of a biological triplicate of three individual wells, each with dozens of cells (n=3). Individual biological replicates are displayed with white circles. (B) Fluorescence-scanned SDS-PAGE gel used to quantify the remaining fluorescent BSA after incubation with *Capsaspora*. The fl-BSA bands were quantified to generate **Figure 4C**.
