## Supporting Information 2 for "Chemokinesis by a microbial predator"

### SII.1 Introduction

Schistosomes release chemicals, once *Capsaspora owczarzaki* cells have attached themselves to the surface. These chemicals have been shown to induce positive chemokinesis in protists, chemokinesis being a chemical dependent change in the locomotion speed of cells. *C.owczarzaki* achieve movement by extension of many thin filopodia which are distributed across a fairly stable actin cortex, which they then use to 'drag' themselves in the target direction which is dependent on adhesion of filopodia to the substrate [1]. Without making any assumption about how the chemokinetic effector acts on the migration mechanism, we assume that positive chemokinesis speeds up *C.owczarzaki* displacement. These protists also seem to display irreversible attachment to schistosomes, as once they pierce the schistosome they are attached until their prey dies. We seek to answer if this chemokinetic response will aid the predator protists, in combination with irreversible attachment and whether this chemokinetic response will aid protists seeking new prey by encouraging dispersal from dead cells.

### SII.2 Model

We will seek to model our problem in a hybrid fashion, modelling the dispersion of schistosome exudates in a continuum and modelling protist migration with an agent based approach.

#### SII.2.1 Chemical governing equations

We will assume a 2-D plane polar coordinate system to model the dispersion of exudates from the schistosome surface, where $c(r, \theta)$ is the concentration of exudates $(mg/ml)$ and $a(\theta)$ is the protist density (cells/$\mu m$) at the surface of the schistosome.

In our geometry $r = 0$ corresponds to the centre of the schistosome, which we will assume is stationary and circular for the sake of modelling simplicity. Although they move very little in in vitro studies and are broadly circular in shape, so this is not a incredulous assumption. Then we have the surface of the schistosome corresponding to $r = r_s$. We assume that $r_s$ is roughly $32\mu m$ based on microscopic measurements. We will then assume that the exudates that are secreted from the schistosome upon predation undergoes isotropic diffusion with constant diffusivity $D_c$, so that the equation of transport of schistosome exudates is given by

$$\frac{\partial c}{\partial t} = D_c \nabla^2 c, \tag{1}$$

where the Laplacian in plane polar coordinates is given by

$$\nabla^2 = \frac{\partial^2}{\partial r^2} + \frac{1}{r}\frac{\partial}{\partial r} + \frac{1}{r^2}\frac{\partial^2}{\partial \theta^2}, \tag{2}$$

with boundary conditions given by

$$\frac{\partial c}{\partial t}(r_s, \theta) = f(a), \quad \frac{\partial c}{\partial t}(L, \theta) = 0,$$

where $L$ is the domain length, with no-flux at the end of edge of the circular domain and $f(a)$ is the function that determines the rate of exudation from the schistosome surface. Since these exudations begin when the surface of the schistosome is pierced by a protist, it would make sense that exudation rate is in some way proportional to protist density at the surface, a sensible form of $f$ would be

$$f(a) = b \tanh\left(\frac{a}{a_0}\right) \tag{3}$$

where $b$ is the exudation rate and $a_0$ is the protist concentration that would lead to the exudation rate being roughly $3b/4$. As it would be reasonable to assume that there is a saturation point where more protists would not lead to greater exudation locally, so a tanh like response would be appropriate.

### SII.2.2 Protist dynamics

To model our protists, we will take an agent based approach, meaning we will model each protist individually, then take the average density at grid points in order to determine the value of equation 3. We will model i protists, with $\boldsymbol{x}_i(t)$ being the location of the centre of the ith protist $(x, y)$ at time $t$. We will assume that each protist has an orientation angle based on which filopodia has been extended to achieve motion, then travels in that straight line direction for a fixed distance. Therefore, each protist will have an orientation angle $\phi_i$ which giving each protists an orientation vector $\boldsymbol{p}_i(t) = (cos(\phi_i), sin(\phi_i))$. Then, if we discretise time into blocks $\Delta t$, the position of each protist is updated by the orientation of the active filopodia and current locomotion speed, such that

$$\boldsymbol{x}_i(t + \Delta t) = \boldsymbol{x}_i(t) + v_i(c)\boldsymbol{p}_i(t)\Delta t \tag{4}$$

until current run length, denoted $l_c$, is greater than or equal to $l_t$ which is a random variable determining length of travel before a reorientation event. Once termination length is determined, a new filopodia will be extended until the cell has travelled $l_t$ in

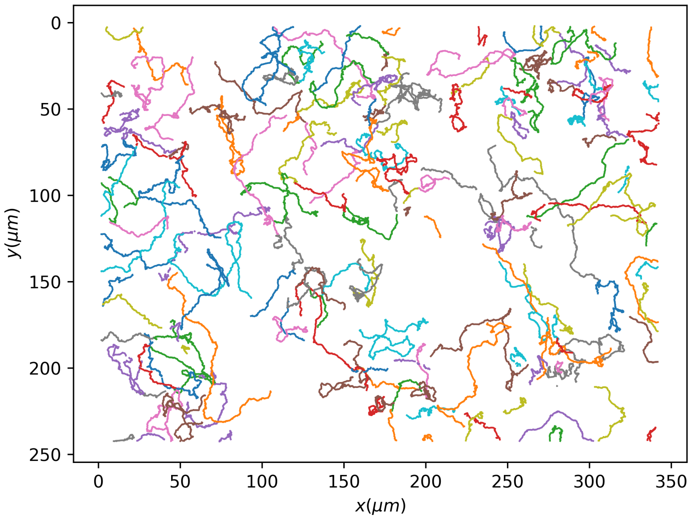

**Fig. SII.1**: Trajectories of WT *C.owczarzaki* cells from Video 1 in [2]

that direction, in other words when $l_c \geq l_t$ the agent will alter their orientation angle according to the equation

$$\phi_i(t + \Delta t) = \phi_i(t) + \Delta\phi \tag{5}$$

where $\Delta\phi$ is a random variable representing the difference between the orientation of the current active filopodia and the next to be extended.

#### SII.2.2.1 Motility statistics

Since the termination length and change in reorientation angle in reality would be dependent on properties of the cell, such as filopodia length, and the change in orientation would be dependent on adhesion of filopodia with the substrate, a physically realistic model would be out of scope of this project. Instead, we opt for a data driven approach, deriving aggregate statistics of *C.owczarzaki* and using those to construct our random walk dynamics.

In order to analyse the motion of *C.owczarzaki*, Video 1 from [2], which displays wild type *C.owczarzaki* cells moving on a homogenous substrate, was used. A median filter was applied, then the motion of the cells over the assay period was analysed using Trackpy [3], the resulting trajectories can be seen in figure SII.1

Then the trajectories were analysed to determine: when cells reorientated, the resultant change in orientation angle and the distance travelled between reorientation events. A kernel density estimate (KDE) of distance between reorientation events $dr$ was fitted using the Seaborn python library [4], which can be seen in SII.2.

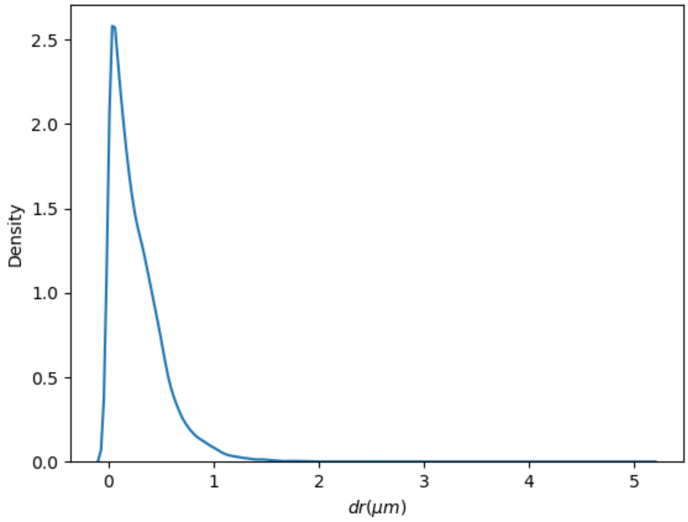


**Fig. SII.2**: KDE of displacement between reorientation events, $\langle dr \rangle = 0.27\mu m$, $Var(dr) = 0.065\mu m$

As we can see, the displacements are exponentially distributed, so it would be reasonable to model $l_t$ as an exponentially distributed random variable with rate $\lambda = 1/\langle dr \rangle = 3.71\mu m$, in other words $l_t \sim \text{Exp}(3.71\mu m)$. In order to determine $\Delta\phi$, we measured the angle between the new and old orientation angles for each cell at each reorientation event. The KDE of $\Delta\phi$ can be seen in figure SII.3. As we can see by observation, the changes in orientation angle are reasonably normally distributed with mean $\langle \Delta\phi \rangle \approx 0$ and truncated between minimum and maximum values $-\pi$ and $\pi$ respectively. Therefore, we can assume that $\Delta\phi$ is a truncated normally distributed random variable, in other words $\Delta\phi \sim \text{TN}(0, 1.84, -\pi, \pi)$. However, for this to be a valid assumption, we will need to look at the autocorrelation of $\Delta\phi$, which is given by

$$R(\tau) = E[\Delta\phi_i \Delta\phi_{i+\tau}], \tag{6}$$

where $i$ is the index of reorientation events and $\tau$ is the lag in that index. The autocorrelation can tell us how much the current orientation change is influenced by previous orientation changes. The autocorrelation can be seen in figure SII.4, as we can see there is a slight negative correlation between the current and last reorientation. Meaning a cell is slightly more likely to choose an orientation to the left (L) or right (R) of its current orientation, if at the last reorientation event it chose an orientation to the R or L of its previous orientation respectively. Although the autocorrelation for $\tau = 1$ is very small it is worth investigating, so we calculated the probability of two consecutive L turns (LL) or R turns (RR) compared to alternating L/R turns (LR/RL), from our trajectories. These probabilities can be seen in figure SII.5, as we can see, after

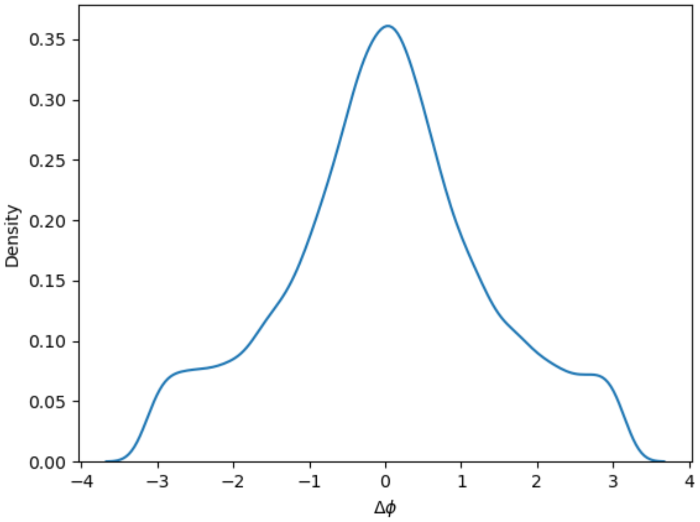


**Fig. SII.3**: KDE of $\Delta\phi$, the difference between new and old orientation angles following a reorientation event, $\langle\Delta\phi\rangle = 0.01$radians, $Var(\Delta\phi) = 1.84$radians

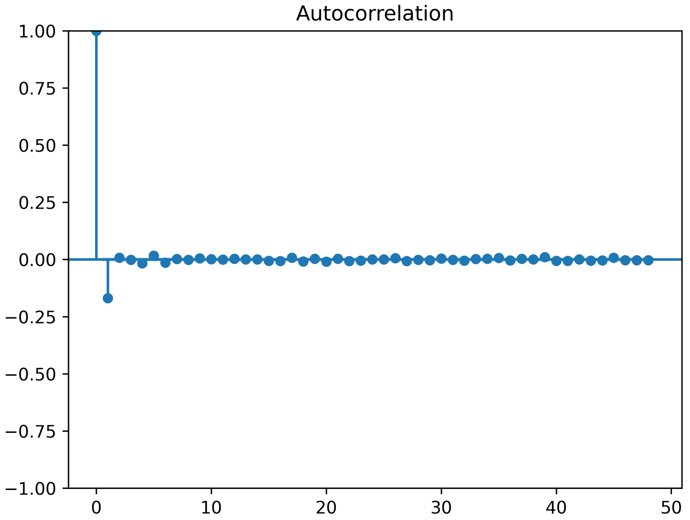


**Fig. SII.4**: Autocorrelation of $\Delta\phi$, x axis indicated lag in terms of number of reorientation events

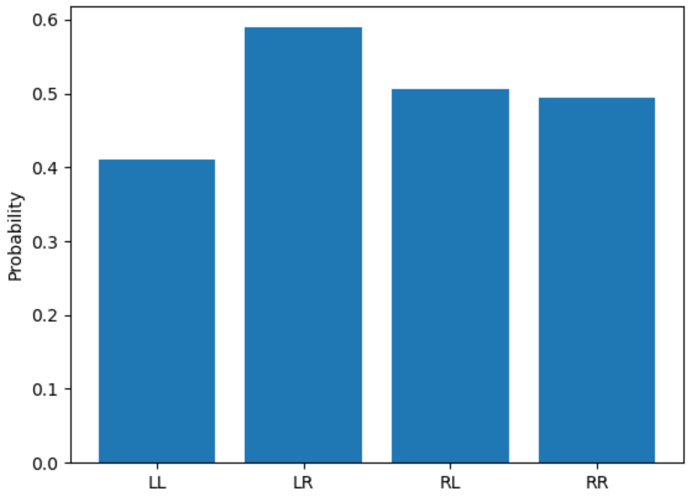


**Fig. SII.5**: Probability of a cell reorientating to the left (L) or right (R) of its current orientation, depending on whether it orientated to the L or R of its previous orientation

a R turn cells are equally likely to turn L or R but after a L turn they slightly more likely to make a R turn. Since this difference is negligible and the autocorrelation is very low, it would not be unreasonable to assume then that reorientation events are independent from each other. Therefore we can sample $\Delta\phi$ directly at each reorientation events and for each cell, and do not have to consider the impact of any previous reorientations of that cell.

Therefore, our data driven approach would see our protist move in a straight line updating its location according to equation 4 until it has traversed $l_t \sim \text{Exp}(3.71\mu m)$. Then it would alter it's orientation angle according to equation 5 where $\Delta\phi \sim \text{TN}(0, 1.84, -\pi, \pi)$.

#### SII.2.2.2 Irreversible attachment

Since protists display irreversible attachment, we will say that when any protists updates their location such that $||x_i(t + \Delta t)|| \leq r_s$, they will become irreversibly attached, at the point where where the proposed trajectory of the protists intersects the circle defining the schistosome ($r = r_s$). We will then define $\eta_i$ as the residence time of the i'th protist at the schistosome surface and use this, alongside the average residence time of all protists

$$\bar{\eta} = \frac{1}{n}\sum_{i=1}^{n}\eta_i \qquad (7)$$

to determine success of predation. That is the say, we will consider the protist population with highest average residency time at the schistosome surface, the more 'successful' predators. As we will need some metric to determine if chemokinesis in combination with irreversible attachment is beneficial in protists predation of schistosomes.

#### SII.2.3 Chemokinesis

We are interested in seeing if chemokinesis in combination with irreversible attachment will benefit protists in their predation of schistosomes. In order to do this we will need to select a functional form of $v(c)$

We will assume that all protists travel at their base speed without chemical bias. Although protists have a distribution of base travel speeds, we will assume they all travel at the population average base speed given by $v_0$. Then we will assume that when $c$ approaches some critical value $c_0$ locomotion speed will increase until saturating at some value $v_c$ above the base speed. This will give us the following expression for a chemokinetic speed curve

$$v(c) = v_0 + v_c \left( \frac{\tanh\left(k\left(c - c_0\right)\right) + \tanh\left(kc_0\right)}{1 + \tanh\left(kc_0\right)} \right) \tag{8}$$

where $k$ is the parameter which determines how steep the transition between $v_0$ and $v_0 + v_c$ is, lower values of $k$ would lead to a more gradual transition and higher values would lead to a sharper more step like transition in response to greater values of $c$.

#### SII.2.4 Nondimensionalisation

We will now nondimensionalise our system of equations, nondimensional quantities will be denoted with an asterisk. Our system can be nondimensionalised using the following scalings,

$$(x^*, y^*, r^*, l_c^*, l_t^*) = \frac{1}{r_s}(x, y, r, l_c, l_t),$$
$$v^* = \frac{v}{v_0}, \quad t^* = \frac{v_0 t}{r_s}, \quad a^* = r_s a, \quad c^* = \frac{c}{c_0},$$

dropping the asterisks for notional clarity, our nondimensional equation of chemical transport is given by

$$\frac{\partial c}{\partial t} = \gamma \nabla^2 c \tag{9}$$

with boundary conditions given by

$$\frac{\partial c}{\partial t}(1, \theta) = \beta \tanh\left(\frac{a}{\alpha}\right), \quad \frac{\partial c}{\partial t}(\tilde{L}, \theta) = 0 \tag{10}$$

and nondimensional parameters

$$\gamma = \frac{D_c}{v_0 r_s}, \quad \tilde{L} = \frac{L}{r_s}, \quad \alpha = r_s a_0, \quad \beta = \frac{r_s b}{v_0 c_0}.$$

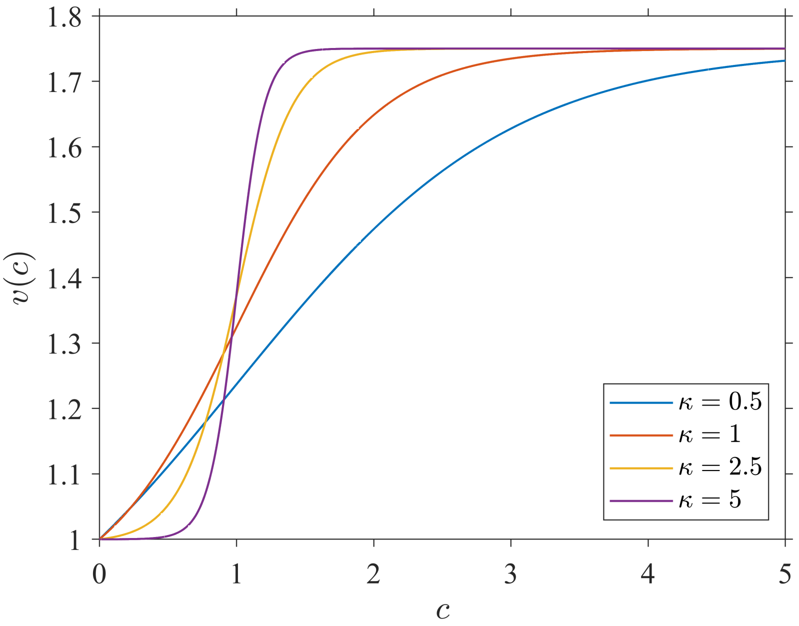


**Fig. SII.6**: Chemokinetic response curves given by equation 11 plotted for a variety of values of $\kappa$, with $\xi = 3/4$

Then, our nondimensional chemokinetic speed function is given by

$$v(c) = 1 + \xi \left( \frac{\tanh(\kappa(c-1)) + \tanh(\kappa)}{1 + \tanh(\kappa)} \right) \tag{11}$$

with nondimensional parameters

$$\xi = \frac{v_c}{v_0}, \quad \kappa = kc_0,$$

where $\kappa = 0$ corresponds to the case of no chemokinesis and the speed curves for a variety of $\kappa$ can be seen in figure SII.6, as we can see higher values of $\kappa$ lead to a much sharper step-like chemokinetic response. The equations of motions for the amoeboid agent remain unchanged in nondimensional form, except with dimensional quantities replaced by their nondimensional counterparts.

### SII.3 Simulation

We will now perform numerical simulations to determine the effect of chemokinesis on residence time of protists, using a range of chemokinetic parameter $\kappa$. We will then repeat these simulations with initial conditions representing several dead schistosomes in the local proximity to assess the effect of this residency times, to see if chemokinesis can act as a dispersion agent away from dead schistosomes, by creating ‘exclusion zones', leading to higher residence times on the living schistosomes. All simulations will be run until $t = 15$, which roughly corresponds to 1 hour based on observed measurements of the timescale.

#### SII.3.1 Numerical details

We will now briefly outline the details behind our numerical simulations, which were carried out in MATLAB.

##### SII.3.1.1 Chemical simulation

The profile for the Chemical was solved on a 2-D polar coordinate grid with the radius discretised into M grid points between 1 and $\tilde{L}$, spaced evenly by h, and the polar coordinates were discretised into M-1 evenly spaced grid points between 0 and $2\pi - \theta_h$ where $\theta_h$ is the grid spacing in the polar grid. Then a fourth order finite difference scheme was implemented to find the derivatives of in the radial and polar directions, with periodic boundary conditions applied at on the polar grid and boundary conditions in radial direction given by equation 10. The system was then discretised in time using a positivity preserving forward Euler scheme,

$$c(t + \Delta t) = \begin{cases} c(t) + \Delta t \dfrac{\partial c(t)}{\partial t}, & \text{if } \dfrac{\partial c(t)}{\partial t} \geq 0 \\ c(t)^2 \left( c(t) - \Delta t \dfrac{\partial c(t)}{\partial t} \right)^{-1}, & \text{if } \dfrac{\partial c(t)}{\partial t} < 0 \end{cases} \tag{12}$$

where the positivity preservation scheme was adapted from Wood and Kojouharov [5]. Positivity preservation is important as we obviously cannot admit a negative concentration.

##### SII.3.1.2 Protist simulation

Equations 4 can be directly computed at each time-step, chemical concentration at protist locations were interpolated from grid points using MATLABs 'Scattered-Interpolant' function via natural neighbour interpolation in order to calculate 11. Termination lengths $l_t$ were computed using the quantile method for the exponential distribution, with non-dimensional rate given by, $\lambda \approx 32/0.27$, such that

$$l_t = -\frac{\log{(1-s)}}{\lambda} \tag{13}$$

where $s \sim U[0, 1)$ and was generated using MATLABs built in random number generator. Once termination lengths were exceeded, orientations were updated according to equation 5, where $\Delta\phi$ was generated using the truncated normal variable generator provided by Botev [6].

#### SII.3.2 Initial conditions

Initially protists are located at a random location in the annulus between circles defined at $r = 1$ and $r = \tilde{L}$ such that

$$\boldsymbol{x}_i(0) = r_0(\cos(\theta_0), \sin(\theta_0)) \tag{14}$$

where $r_0 \sim U[1, \tilde{L}]$ and $\theta_0 \sim U[0, 2\pi]$. Then we have two different cases for chemical concentration, in the first case there is no background chemical and chemical concentration only deviates from zero due to schistosome exudations and

$$c(r, \theta, 0) = 0. \tag{15}$$

In the other scenario, there are 4 recently dead schistosomes located at $r = r_c$ and $\theta = \pi/4, 3\pi/4, 5\pi/4, 7\pi/4$, which creates exponential profiles at those locations, giving us initial conditions

$$c(r, \theta, 0) = c_\infty \sum_{i=0,1,2,3} \exp\left(-\frac{\mu(i)}{\sigma_c}\right) \tag{16}$$

$$\mu(i) = \left(\left(r\cos(\theta) - r_c\cos\left(\frac{\pi}{4}(1+2i)\right)\right)^2 + \left(r\sin(\theta) - r_c\sin\left(\frac{\pi}{4}(1+2i)\right)\right)^2\right)^{1/2} \tag{17}$$

where $c_\infty$ is concentration of most dense part of the dead schistosome and $\sigma_c$ determines how sharp the peak of chemical from the dead schistosome is.

#### SII.3.3 Chemokinetic shape parameter effect on residency times

To assess residency times, we then ran our simulations for both initial conditions and various chemokinetic parameters, a snapshot of a simulation with initial conditions from equation 16 and $\kappa = 2.5$ can be seen in figure SII.7.

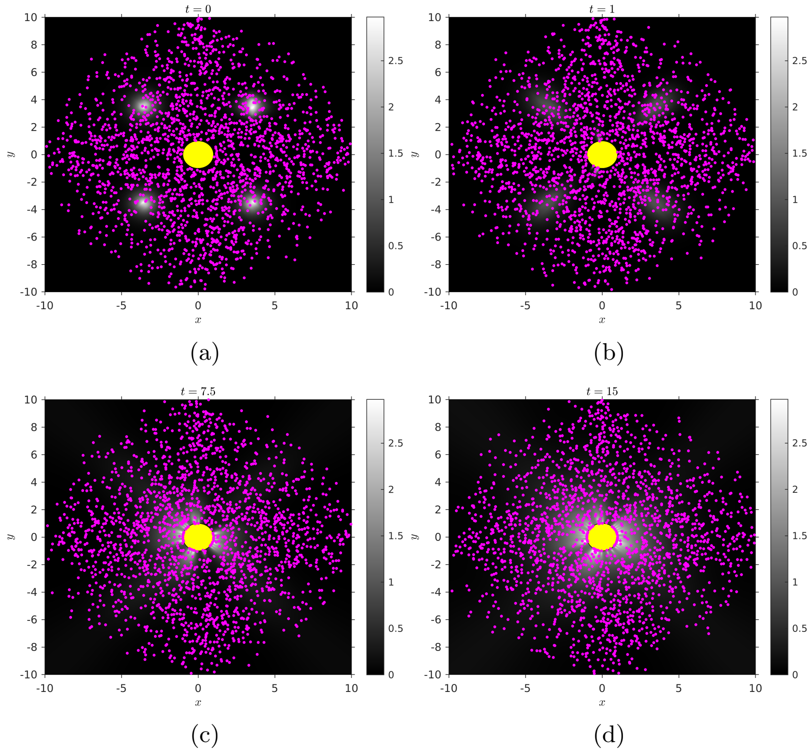


**Fig. SII.7**: Snapshot of simulation of chemokinetic protists (purple dots) in chemical concentration field c indicated by colour bar, predating schistosome (yellow circle) with initial conditions specified in equation 16. Simulation parameters used: $N = 2000$, $\xi = 3/4$, $\tilde{L} = 9$, $\gamma = 0.5$, $r_c = 5$, $\sigma_c = 0.5$, $h = 0.1$, $\Delta t = 10^{-3}$, $c_\infty = 3.5$, $\alpha = 4$, $\beta = 120$, $\kappa = 3.5$.

We can see the average residency times in figure SII.8a, we can see that certain values of our chemokinetic parameter lead to an advantage in residency times. However this relationship is non-linear and it seems that the specific value of $\kappa$, which defines the cells chemokinetic response, profoundly impacts whether or not chemokinesis and irreversible attachment is advantageous or disadvantageous to the predation of schistosomes by *C.owczarzaki*. Although, it is important to note that even in the cases where chemokinetic cells are 'outcompeted' by non chemokinetic cells ($\kappa = 0$), the disadvantage is relatively minor compared to the scenarios in which chemokinesis is conferring an advantage. Therefore, while we can't say whether chemokinesis is advantageous or not without parametrising a specific chemokinetic response, we can say on average a chemokinetic response has higher potential upsides than downsides.

In the case of dead cells acting as a 'dispersal agent', we can see in figure SII.8b that it is entirely dependent on $\kappa$ whether the presence of recently dead schistosomes would be advantageous or deleterious to *C.owczarzaki* cells. Although we can say that for majority of simulated cases, it is advantageous to have these dead cells present. This is most likely due to the fact that cells, in these 'exclusion zones', effectively cover more area as cells will exit the vicinity much faster and are therefore more likely to encounter the schistosome earlier, as the effective space to search is reduced. Although, this depends on which directions cells are facing initially, as we have shown *C.owczarzaki* displays strong directional persistence. So if cells were on average facing towards the schistosome, chemokinesis would most likely aid in predation, and if they were on average facing away from the schistosome, chemokinesis would like inhibit predation. So the stochastic initial conditions would strongly influence the effect of chemokinesis in this scenario, explaining why figure SII.8b does not always show an advantage or disadvantage due to these initial conditions. Although it would be fair to say there is some relationship between $\kappa$ and the presence of these 'exclusion zones', in successful predation by *C.owczarzaki* cells.

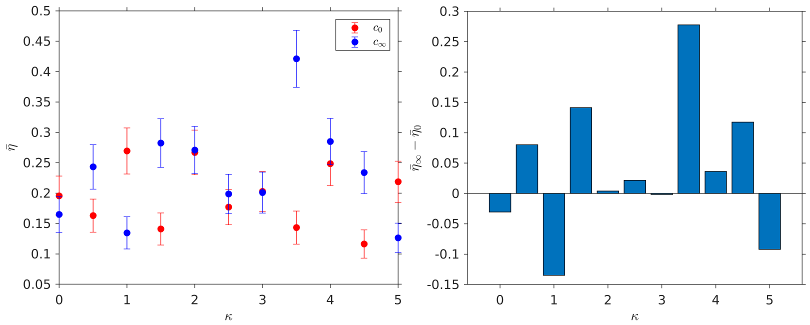


(a) (b)

**Fig. SII.8**: Effect of chemokinetic parameter $\kappa$, on average residency times $\bar{\eta}$, 2 different initial conditions, $c_0$ corresponds to 0 concentration and $c_\infty$ corresponds to equation 16 : (a) $\bar{\eta}$ plotted against $\kappa$, error bars represent standard error $(\pm\sigma(\eta)/\sqrt{N})$ ; (b) Bar chart showing the average residency time for initial conditions in 16 $\bar{\eta}_\infty$ minus average residency times for initial conditions with initially 0 concentration $\bar{\eta}_0$ for the values of $\kappa$ in (a). Simulation parameters used: $N = 2000$, $\xi = 3/4$, $\tilde{L} = 9$, $\gamma = 0.5$, $r_c = 5$, $\sigma_c = 0.5$, $h = 0.1$, $\Delta t = 10^{-3}$, $c_\infty = 3.5$, $\alpha = 4$, $\beta = 120$.

#### SII.3.4 Main text parameters

In the main text, the dose-dependent speeds were fitted to the chemokinetic curve in order to determine the values of $\kappa$ and $\xi$ that corresponds to Bovine Serum Albumen and Human Serum Albumen. These values was found to be $\kappa = 2.5359$, for both BSA and HSA which was used for all simulations in the main text. We then found that BSA corresponds to $\xi = 3/4$ and HSA corresponds to $\xi = 1/2$. The diffusivity of both BSA and HSA were assumed to be the same, which was taken from [7], corresponding to a value $\gamma \approx 13.5$, which was used in all simulations in the main text. As you can see from the main text, this high diffusivity changes the dynamics considerably and leads to a more distinct effect of chemokinesis.

| Parameter | BSA | HSA |
|---|---|---|
| $N$(Number of protists) | 1000 | 1000 |
| $\tau_{max}$(Simulation time) | 2.5 | 2.5 |
| $\alpha$ | 8 | 8 |
| $\beta$ | 50 | 50 |
| $\xi$ | 3/4 | 1/2 |
| $\gamma$ | 13.5 | 13.5 |
| $\kappa$ | 2.359 | 2.359 |
| $\sigma_c$ | 0.5 | 0.5 |
| $r_c$ | 5 | 5 |
| $c_\infty$ | 3.5 | 3.5 |
| $h$ | 0.1 | 0.1 |
| $\Delta t$ | $10^{-4}$ | $10^{-4}$ |

**Table SII.1**: Table of parameters used in simulations of protists responding to schistosomes exuding either BSA or HSA
